# The complete genome of the KOLF2.1J reference iPSC line

**DOI:** 10.64898/2026.03.06.710144

**Authors:** Pilar Alvarez Jerez, Arang Rhie, Juhyun Kim, Prajna Hebbar, Sagorika Nag, Dmitry Antipov, Sergey Koren, Erika Lara, Alexandra Beilina, Nancy F. Hansen, Charles Arber, Jimena Zulueta, Peter Wild Crea, Dhairya Patel, Glenn Hickey, Brian Waltz, Laksh Malik, William C Skarnes, Xylena Reed, Rylee Genner, Kensuke Daida, Caroline B. Pantazis, Francis Grenn, Mike A. Nalls, Kimberley Billingsley, Valentina Fossati, Selina Wray, Michael Ward, Mina Ryten, Andrew B. Singleton, Mark R Cookson, Miten Jain, Benedict Paten, Adam M Phillippy, Cornelis Blauwendraat

## Abstract

While induced pluripotent stem cells (iPSCs) have gained popularity in studying neurodegenerative diseases, the heterogeneity of stem cells used across studies impacts cross-study comparison. The iPSC Neurodegenerative Disease Initiative (iNDI) selected the KOLF2.1J cell line and prioritized its use as a reference standard for studying the effects of pathogenic variants on cell biology due to its stability and neutral neurodegenerative disease genetic risk. This cell line, and its derivatives expressing over 100 variants related to Alzheimer’s disease, Parkinson’s disease, and other neurological diseases, are available for academic and industry access. Current genomic data analyses are limited by the use of a human reference genome that does not capture the complete genetic background of a given iPSC line. While in the future this issue may be partially mitigated by the creation of a comprehensive human pangenome, previous work has shown that generating custom genomes is of value both to characterize the variation present and to serve as a more appropriate genomic reference. Here, we generated and characterized a custom complete genome assembly from KOLF2.1J. Mapping of sequencing reads to a personalized diploid assembly results in more comprehensive mapping compared to traditional linear references (i.e GRCh38). In addition, we provide a comprehensive custom gene annotation along with isoform expression and differential methylation analyses across multiple cell types. The assembly and all additional data is browsable and publicly available. This resource will enable more accurate investigation of the KOLF2.1J cell line and any genomics data generated compared to using traditional generalized references, while also serving as a foundational approach for establishing custom reference assemblies for other high-value iPSC lines.

## Introduction

Although both causal and risk variants have been identified for many neurological diseases, their effects and mechanisms of action are still largely unknown. In response, induced pluripotent stem cell (iPSC) models have gained popularity over recent years as a method to model and explore the role of disease variants on biological processes (Sullivan and Young-Pearse 2017). Additionally, differentiation of iPSCs into disease-relevant cell types, such as neurons and microglia, and more recently organoids, allows for study into cell-type-specific variant effects (Wang et al. 2017; Abud et al. 2017). However, although iPSC usage is gaining popularity and models are improving, results become difficult to compare across labs studying the same disease with these models because of a lack of common reference cell line.

In order to address this limitation, the iPSC Neurodegenerative Disease Initiative (iNDI) identified the KOLF2.1J cell line and prioritized its use as a reference standard for studying neurodegenerative diseases (NDDs) due to its adequate growth properties, stability, and neutral genetic neurodegeneration risk (Pantazis et al. 2022). This line, and its derivatives harboring over 100 variants known to be associated with NDDs, is currently available at The Jackson Laboratory and accessible by academia and industry (https://www.jax.org/jax-mice-and-services/ipsc/cells-collection). This provides labs the ability to use the same reference line, and as such be able to compare results more accurately.

In addition to iPSC variation, current genomic analyses are limited by the use of general reference genomes for sequence alignments (Taylor et al. 2024). While a general reference genome can be an accurate method for detecting single nucleotide variants, reference bias can occur when mapping both genomic and transcriptomic data. A personalized assembly is an improved method to search for large genomic structural variants (Kolmogorov et al. 2023), but generating a high quality personalized assembly requires sequencing data of high depth and long read lengths. Recent sequencing and analysis tool advances have made data generation and genome assembly more feasible. For example, the Verkko assembler is able to automatically assemble and phase entire human chromosomes by combining Pacific Biosciences’s highly accurate HiFi sequencing reads and Oxford Nanopore’s ultra long sequencing reads(Rautiainen et al. 2023)(Jain et al. 2018; Wenger et al. 2019). Another important assembler is Hifiasm (Cheng et al. 2021, 2024). While it was originally created to assemble only from HiFi reads, new developments now allow it to incorporate Nanopore’s ultra long reads for more complete assemblies.

While the process of generating all the data necessary for an accurate personalized assembly can be costly, investing in custom genome references is worth it for a resource that will be used by labs around the world as they are of high value both to characterize the genetic variation present and to serve as a more appropriate reference for downstream functional analyses (Rozowsky et al. 2023). Recent work on the human pangenome has demonstrated substantial gains in variant call accuracy when using personalized references (Liao et al. 2023; Sirén et al. 2024). High-quality custom references have been generated for other cell lines of interest, including HG002 (Hansen et al. 2025), RPE-1 (Volpe et al. 2025), BJ and IMR-90 (Ranallo-Benavidez et al. 2025) allowing for high-precision expression and epigenetic analyses without the issue of reference bias. However, a custom reference does not yet exist for KOLF2.1J.

Recent efforts to assemble a complete “telomere-to-telomere” (T2T) human genome has resulted in the T2T-CHM13 reference genome. This assembly introduced over 200 Mb of new human genome sequence, harboring 1,956 novel gene predictions (Nurk et al. 2022). Here, we present a complete T2T reference genome of the KOLF2.1J cell line to enable more accurate investigation of NDD genomics. Through this process, we also provide a full structural variant characterization of KOLF2.1J, the methylation patterns of multiple differentiated cell types, and annotation of novel RNA isoforms. The KOLF2.1J assembly, annotation, and sequencing data is browsable and all publicly available as a resource.

## Results

### Sequencing statistics of generated KOLF2.1J data

To comprehensively characterize the genomic, epigenetic and transcriptomic landscape of KOLF2.1J and its derived neural cell types, we differentiated multiple cell types from this line, including NGN2-induced neurons, cortical neurons, astrocytes , and microglia, oligodendrocyte lineage cells, and generated diverse datasets using a range of sequencing technologies. Some of these data was used for the assembly construction, while others were used for genome annotation and investigation into the haplotype-specific transcriptome and methylation profiles (Figure 1). This strategy utilized both short- and long-read technologies including ONT, PacBio, Illumina, CAGE-seq, and Hi-C to generate the necessary data. Table 1 summarizes the sequencing statistics for the sequencing runs performed.

**Figure 1:**
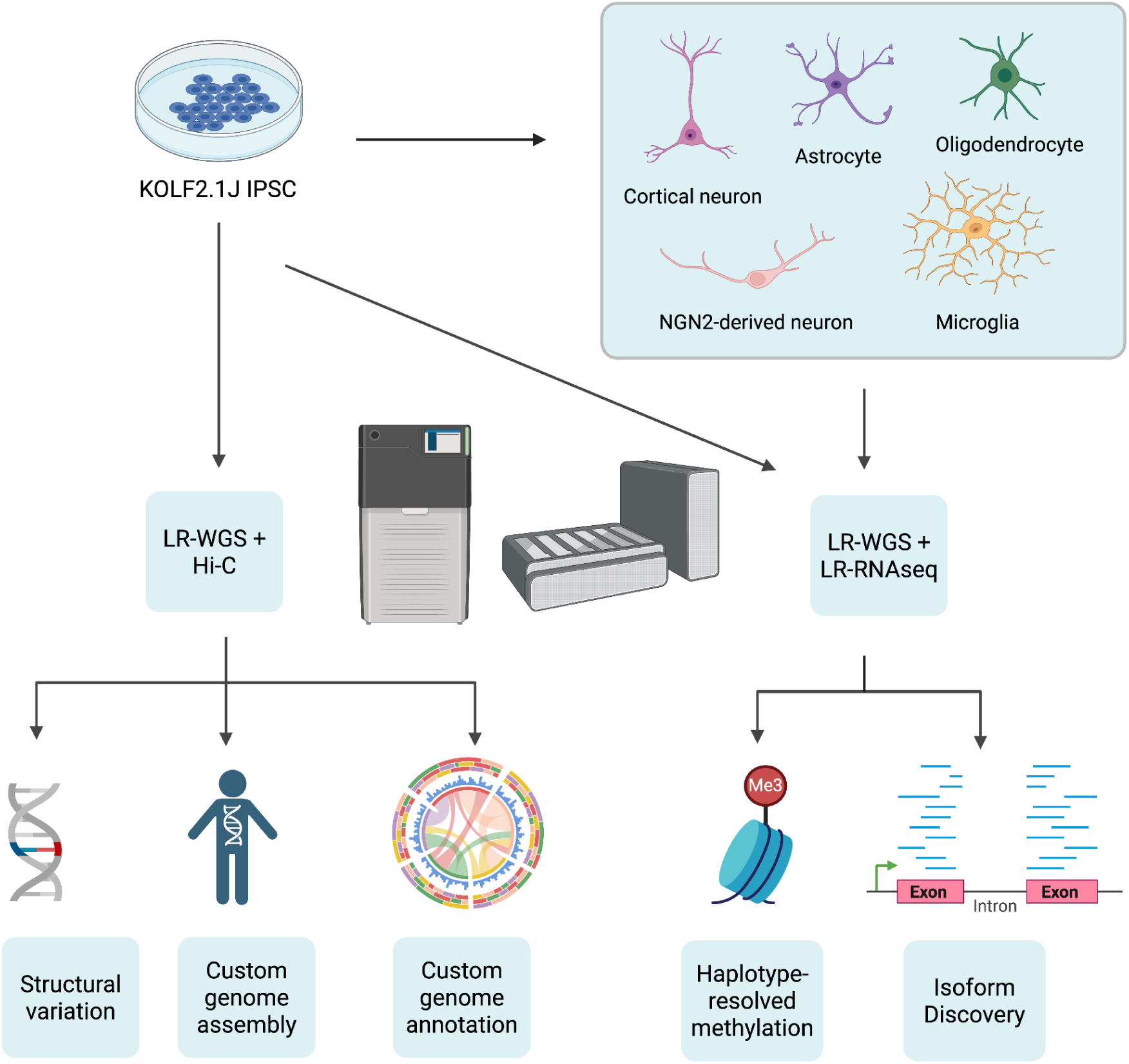
Overview of data generation and analysis. (Left) Oxford Nanopore Technologies (ONT) ultra-long whole genome sequencing (WGS), Pacific Biosciences (PacBio) WGS, and Hi-C data were generated from the KOLF2.1J parental cell line in its iPSC state to generate a custom genome assembly and annotation. The sequencing data was additionally used to catalog structural variants in our line. (Right) KOLF2.1J IPSC was differentiated into multiple cell states: astrocytes, cortical neurons, NGN2 derived neurons, microglia, and oligodendrocytes. We then generated ONT WGS data as well as ONT and PacBio RNA sequencing data to explore methylation and transcript differential expression across cell states and haplotypes. Created with biorender.com

**Figure 2:**
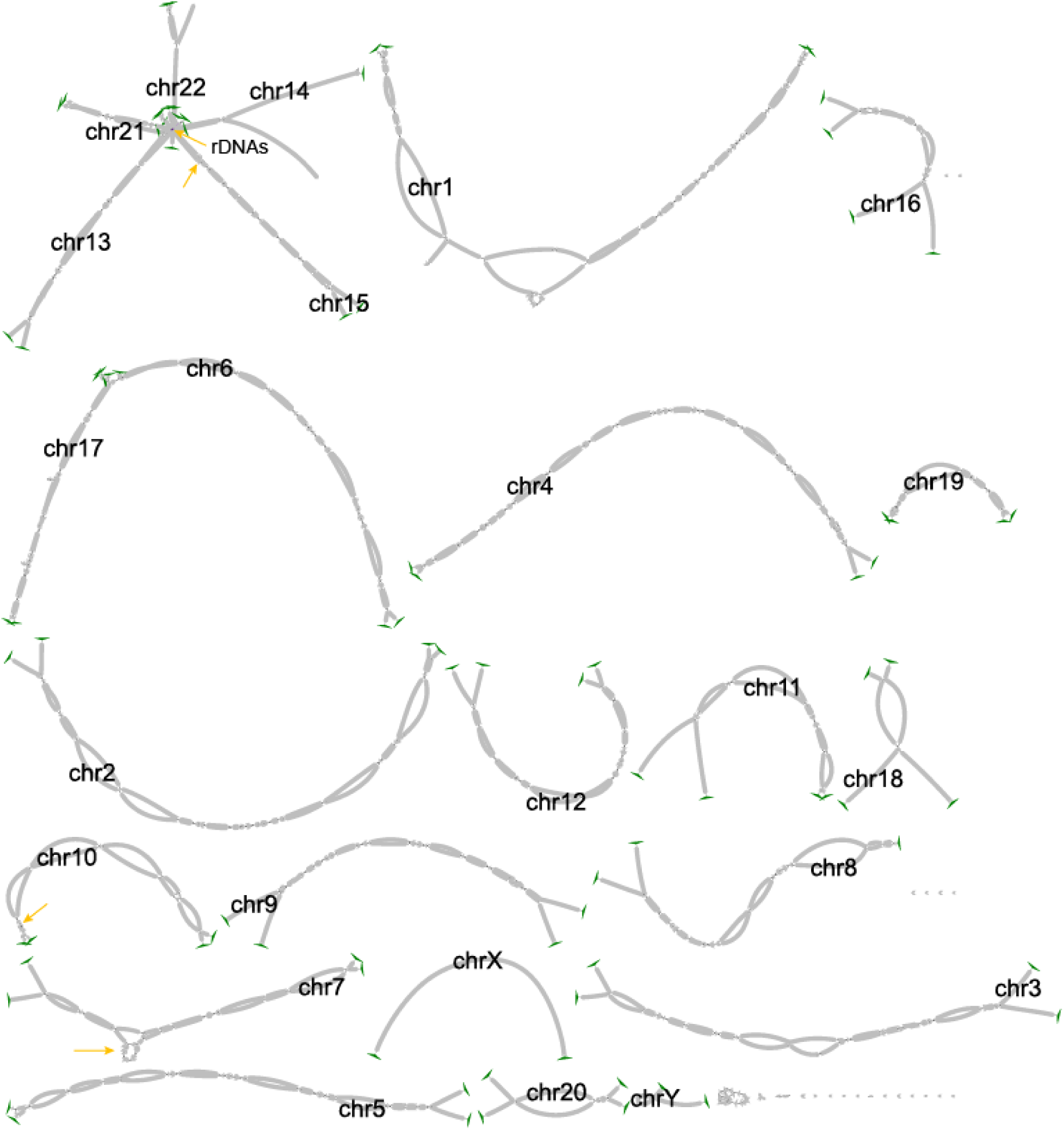
Bandage plot of the KOLF2.1J assembly. The yellow arrows point to unresolved tangles left as gaps in final KOLF2.1J assembly. Green triangles denote telomere motifs found and the end of the last node per chromosome haplotype.

**Table 1:** Summary of sequencing statistics across technologies. N50 = Average read length, Kb = Kilobases, Gb = Gigabases, FC = number of flowcells, ONT = Oxford Nanopore Technologies, PacBio = Pacific Biosciences, HiFi = High Fidelity, UCL = University College of London, NYSCF = New York Stem Cell Foundation

| Sequencing Type | Sequencing Technology | Technology Subtype | Cell Type | N50 (Kb) | Data (Gb) | # of FC |
| --- | --- | --- | --- | --- | --- | --- |
| DNA | ONT | Ultra Long (R9) | iPSC | 91 | 126.3 | 19 |
|  |  | Standard (R9) | iPSC | 32.8 | 109.34 | 1 |
|  |  | Standard (R9) | Microglia | 28.5 | 258.2 | 5 |
|  |  | Standard (R9) | Cortical neuron | 27.1 | 227.6 | 5 |
|  |  | Standard (R9) | NGN2 neuron | 29 | 216.4 | 4 |
|  |  | Standard (R9) | Astrocyte (UCL) | 22.2 | 112 | 1 |
|  |  | Standard (R9) | Astrocyte (NYSCF) | 24.4 | 66.4 | 1 |
|  | PacBio | HiFi | iPSC | 18.8 | 185.3 | 5 |
|  | Illumina | NovaSeq 6000 S4 | iPSC | 0.151 | 118.7 | 1 |
| RNA | ONT | Standard (R9) | iPSC | 1.2 | 87.2 | 1 |
|  |  | Standard (R9) | Microglia | 1.2 | 54.5 | 1 |
|  |  | Standard (R9) | Cortical neuron | 1.5 | 84.1 | 1 |
|  |  | Standard (R9) | NGN2 neuron | 1.1 | 29 | 1 |
|  |  | Standard (R9) | Astrocyte (UCL) | 1.1 | 55.4 | 4 |
|  |  | Standard (R9) | Astrocyte (NYSCF) | 1.3 | 55.3 | 1 |
|  |  | Standard (R9) | Oligodendrocyte | 1.1 | 16.74 | 2 |
|  | PacBio | Kinnex | Microglia | 1.81 | 133.9 | 3 |
|  | Illumina | NovaSeq X Plus 25B | iPSC | 0.151 | 19.4 | 1 |
|  |  | NovaSeq X Plus 25B | Cortical neuron | 0.151 | 17.7 | 1 |
| **CAGEseq** | DNAForm | NA | iPSC | 0.075 | 3 | 1 |
|  |  | NA | Cortical neuron | 0.075 | 3.2 | 1 |
|  |  | NA | Microglia | 0.075 | 3.1 | 1 |
| **Hi-C** | Illumina | NA | iPSC | 0.150 | 187.8 | 1 |
|  | Illumina | NA | Microglia | 0.150 | 208.8 | 1 |

### Assembly Statistics of the KOLF2.1J assembly

We constructed the KOLF2.1J assembly with Verkko v1.4 (Rautiainen et al. 2023) using the HiFi, ONT-UL reads and Hi-C reads from the iPSC cell state. The assembly was polished with HiFi, ONT and Illumina reads following the T2T-Polish recipe (Mc Cartney et al. 2022). In parallel, the assembly graph was manually curated and patched for the remaining gaps with an updated version of Verkko, v2.0 (Antipov et al. 2025). After polishing and curation, the Phred-scaled consensus quality (QV) score improved from Q63.5 to Q67.4 measured with HiFi and Illumina hybrid 31-mers (Rhie et al. 2020). All chromosomes were resolved from telomere-to-telomere, except for the 10 rDNA loci in the short-arms of the acrocentric chromosomes and 4 outside the rDNAs locus (Supplementary Table 1). A 1 Mbp sized gap was placed to represent the missing rDNA arrays. Coverage based analysis indicated potential remaining issues (265.4 kb in total) in addition to the collapsed rDNAs flanking the rDNA gaps (1.9 Mbp in total, Supplementary Table 1). The primary haplotype for each chromosome (hap1) was chosen based on higher QV and fewer gaps, to ensure greater contiguity and higher base-level quality (average Q70.6). The Y chromosome was included in the assembly construction and polishing steps, however it was subsequently removed to abide with the KOLF2.1J sharing consent policy.

### Structural Variant Calling of the KOLF2.1J genome

To explore and catalog SVs in KOLF2.1J, we ran hapdiff, an SV calling package for diploid assemblies. After filtering structural variants for a minimum length of 50bp and removing centromere and satellite regions designated by Genome In A Bottle (GIAB), we were left with 25,521 total SVs out of which ∼16,000 were called as insertions deletions and ∼9,400 called as deletions. There is a known insertion bias in GRCh38 that has previously been reported due to missing sequences or collapsed tandem repeats, yet our numbers are in line with the bias seen in the T2T-CHM13 paper (Aganezov et al. 2022). Both haplotypes had a similar number of total SVs, around ∼17,000 for each (counting both homozygous and heterozygous variants). After annotation with ANNOVAR, we looked specifically at coding variants by filtering for those annotated as “exonic” or “splicing”, given that these are likely having the largest effect on gene expression. This resulted in 188 SVs in KOLF2.1J that were overlapping with a potentially coding region of the genome (Supplementary Table 2). We then manually checked these potentially exonic disrupting SVs to look at their effect on RNA expression (mapped to hg38). Two examples on how the SVs align with changes in RNA are shown in Figure 3.

**Figure 3:**
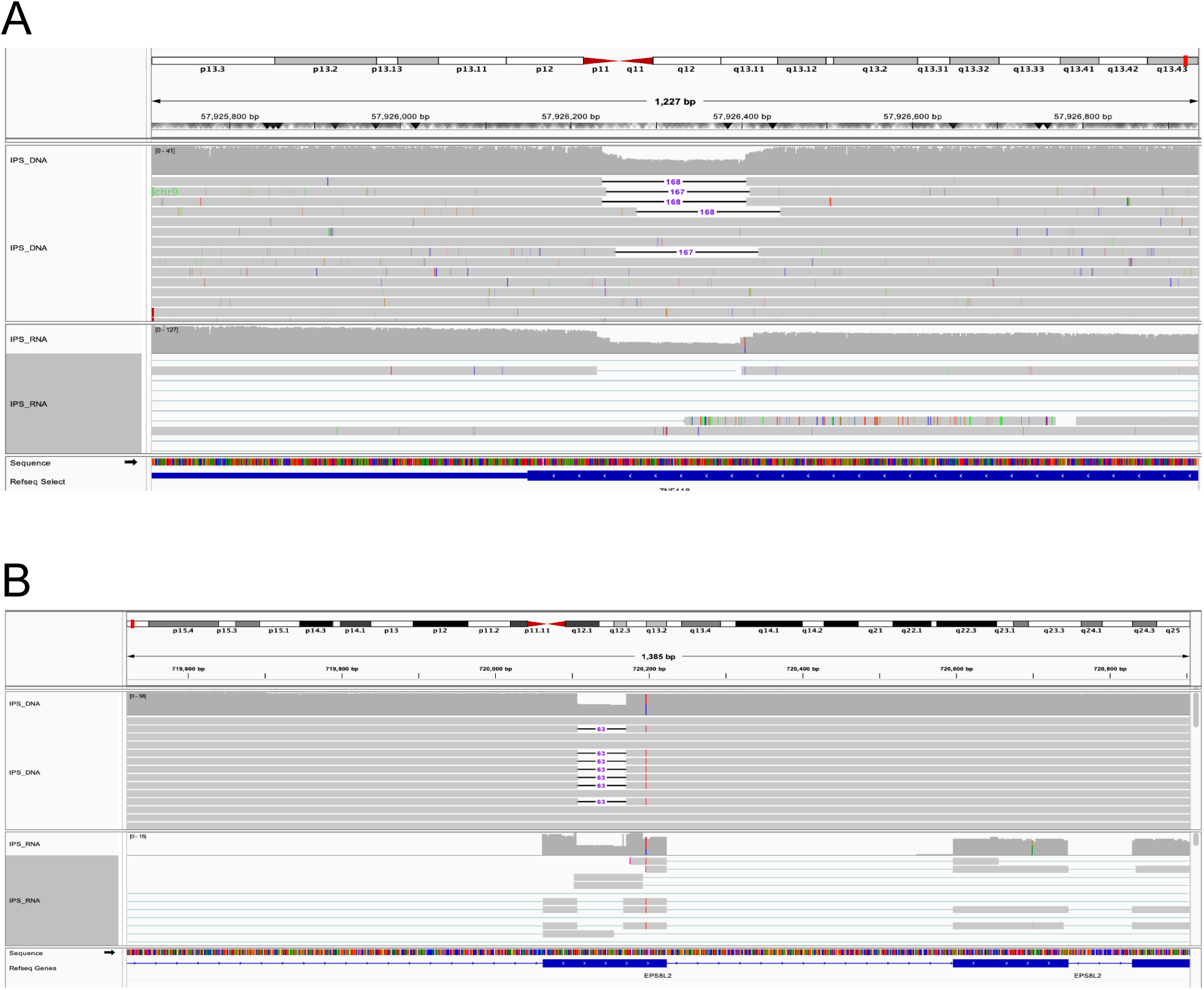
Alignment of coding SVs with matching RNA data on IGV. A) The DNA track at the top shows a heterozygous ∼167bp deletion overlapping exon 4 in the *ZNF418* gene, located in chr19. The matching RNA track on the bottom shows a reduction of approximately half in RNA expression in line with the SV region B) On top, a 63bp heterozygous deletion in exon 5 of the *EPS8L2* gene resulted in a reduction of RNA shown in the bottom track.

### Custom Gene Annotation of the KOLF2.1J Genome

We generated gene annotations for each haplotype of the KOLF2.1J assembly using the Comparative Annotation Toolkit (CAT) using the transMap, liftoff, and miniprot modes, to map genes from GENCODE v47. In addition to these, we mapped the ONT long read RNA sequencing data from five different cell types derived from KOLF and ran CAT with the AUGUSTUS ab initio mode to annotate novel isoforms of genes. The annotation set contains 81,016 genes on hap1 (autosomes + chrX) and 78,642 genes on hap2 (autosomes + chrY) (Table 2). Using the same GENCODE set to annotate genes on CHM13, 81,544 genes were annotated on the CHM13 autosomes + chrX and 79,370 genes on CHM13 autosomes + chrY. All single copy genes annotated in GENCODE were successfully lifted over to the 2 KOLF2.1J haplotypes with all the differences coming from copy number variable genes. A very small number of protein coding gene transcripts have frameshifting variants in KOLF2.1J and a majority (68%) of these belong to gene families like GAGE, MAGE, etc. that contain variable number tandem repeats (VNTRs) (Table 2). There are a total of 201 genes that are CNV between KOLF2.1J hap1 and CHM13 & 256 between KOLF2.1J hap2 & CHM13. This list includes members of large neurodevelopmental gene families like NBPF, NPIP & TBC1D3, and known CNV gene families like GOLGA, SPDYE and AMY.

**Table 2:** Overview of KOLF2.1J custom genome annotation. Overview of number of genes found in each KOLF2.1J haplotype compared to the CHM13-T2T and GrCh38 references.

|  | KOLF2.1J_<br>HAP1<br>(1-22+X) | KOLF2.1J_<br>_HAP2<br>(1-22+Y) | CHM13<br>(1-22+X) | CHM13<br>(1-22+Y) | GRCh38<br>(1-22+X) | GRCh38<br>(1-22+Y) |
| --- | --- | --- | --- | --- | --- | --- |
| **Genes** | 81,016 | 78,642 | 81,544 | 79,370 | 78,014 | 75,731 |
| **Protein Coding Genes** | 20,119 | 19,343 | 20,125 | 19,348 | 20,035 | 19,230 |
| **Frameshifted Transcripts** | 107 | 94 | - | - | - | - |
| **Novel Protein Coding**<br>**Isoforms** | 1878 | 1619 | - | - | - | - |

With this complete gene annotation dataset, we confirmed two previously reported CNVs in coding regions. The first is a heterozygous CNV in chr6p22 resulting in hap1 of the KOLF2.1J assembly having a truncated copy of the *JARID2* gene & missing the *DTNBP1* gene. The second is the truncation of the ASTN2 gene & deletion of its antisense-gene in the paternal haplotype due to the chr9q33 CNV. (Gracia-Diaz et al 2024)

The KOLF2.1J annotation also contains 1878 & 1619 novel predicted protein coding isoforms across the two haplotypes, identified using the IsoSeq data. Novel isoforms have been identified in genes like *PRKAR1B*, a promising candidate gene for neurodevelopmental disorders (Marbach et al 2021) and *DCTN1*, a gene essential for intracellular transport within neurons.

### Improvements in mapping using a custom genome

Read mapping is a fundamental step in genomic data analysis, enabling critical downstream applications such as variant calling and expression quantification. However, mapping to a generalized reference, like GRCh38, containing underlying differences to the sample being mapped introduces reference bias. This is due to easier mapping for reads that match the reference and difficulties to map those that do not (Lin et al. 2024)(Degner et al. 2009). Consequently, read mismapping can introduce spurious or missed variant calls or skew gene expression estimates. To address this, the genomics community is increasingly adopting more complete assemblies such as the T2T-CHM13, which resolved previously incomplete regions (Nurk et al. 2022) (Schneider et al. 2017). Additionally, the Human Pangenome Reference Consortium (HPRC) is developing a pangenome comprising high-quality reference haplotypes from diverse populations integrated into a genome graph that acts as a more comprehensive substrate for mapping (Liao et al. 2023). Building on this resource, personalized pangenome references, which condense a larger pangenome down to (ideally) just the variation relevant to a given sample, provide an important way to eliminate variation that is irrelevant to a given sample being mapped (Sirén et al. 2024). However, a population derived pangenome will not include all the rare and private variants present in a new sample unless that sample is itself present within it. Here, we sought to characterize differences in mapping across multiple reference representations and to directly compare to two representations of the KOLF2.1J assembly, thereby evaluating the advantages of using a customized genome for downstream mapping-based analyses in genomics research.

In what follows, the KOLF2.1J diploid assembly is a linear reference in which the maternal and paternal haplotypes are represented as separate sequences. In contrast, the KOLF2.1J diploid graph is a graph-based reference in which shared regions of the two haplotypes are merged, and divergence is represented only at haplotype-specific variant sites.

Short-read mapping statistics revealed that although gross alignment rates were uniformly high (∼98–99%) for KOLF2.1J short read Illumina WGS data, both the diploid graph and diploid assembly captured ∼13% more perfectly mapped reads (reads mapped without any differences) than GRCh38 (Figure 4B, Supplementary Table 3). The fraction of perfectly mapped reads follows a consistent performance hierarchy that broadly agrees with what is expected: Diploid Graph ∼ Diploid Assembly > Personalized Pangenome > Pangenome > T2T-CHM13 > GRCh38. Additional metrics including gaplessly-mapped read counts (reads mapped without insertion or deletions), mean alignment scores, and error rates (insertions, deletions, mismatches, and soft-clips) reinforced this pattern, with the diploid graph and diploid assembly consistently outperforming other references (Figure 4B,C). We did notice a bias towards deletion calls which we believe is due to the higher difficulty of calling insertions from short read data. While none of these statistics prove the correctness of mapping, the overall trends are supportive of improved empirical mapping, particularly relative to existing linear references, and support earlier simulation studies and evaluations of variant calling to pangenomes relative to linear references (Aganezov et al. 2022)(Sirén et al. 2021)(Garrison et al. 2018).

**Figure 4.**
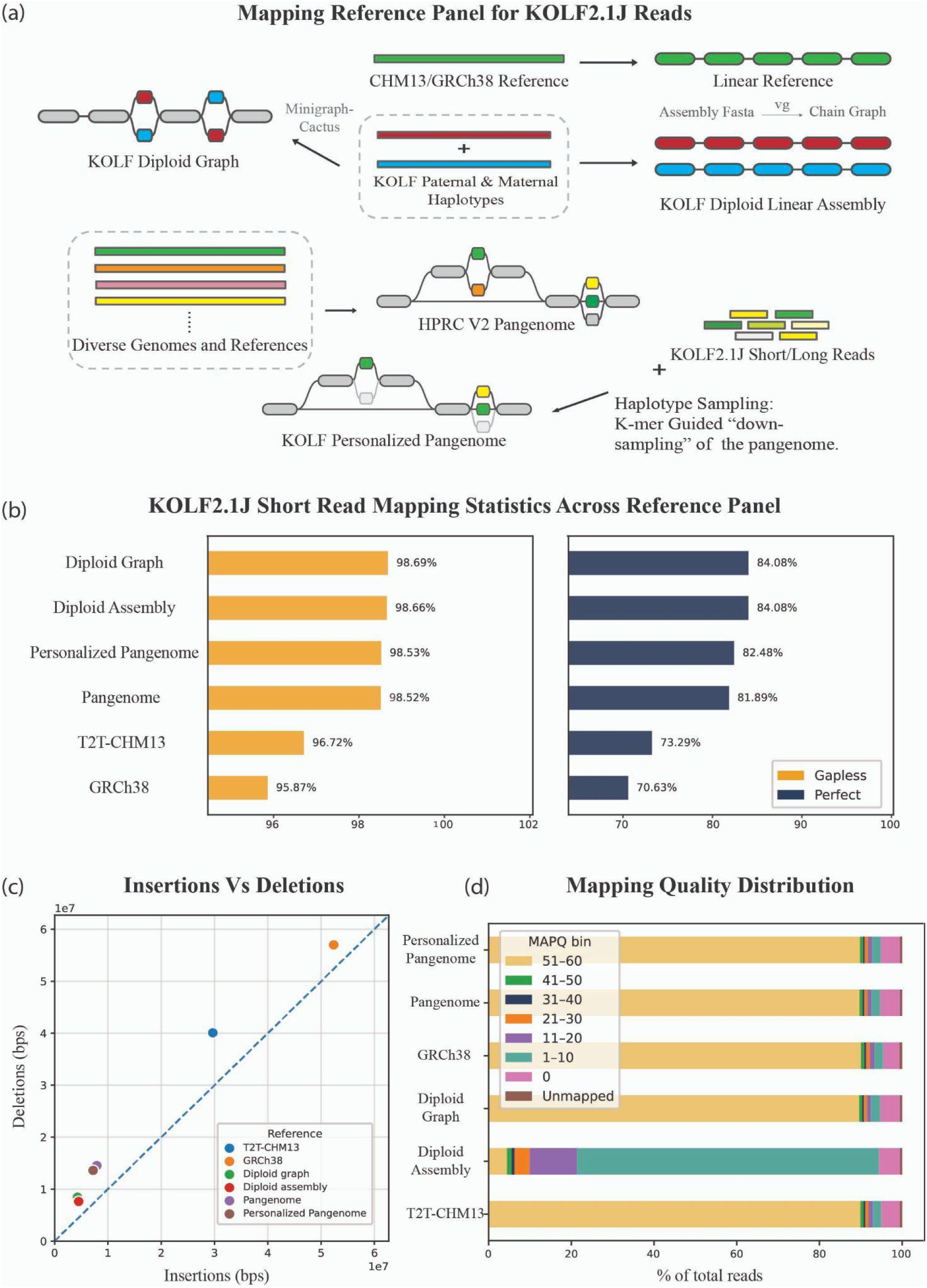
Mapping performance evaluation of KOLF2.1J Illumina short-read data across multiple reference genomes. (a) Mapping Reference Panel for KOLF2.1J Reads. (b) KOLF2.1J short read mapping statistics across the reference panel, representing the total aligned, total perfect and total gapless (soft clips included) reads as a fraction of total reads. (c) Insertions Vs Deletions, plot comparing the total number of insertions versus deletions (measured in base pairs) detected for each reference. (d) Mapping quality distribution, displaying the fraction of reads assigned to different MAPQ bins.

However, mapping quality (MAPQ) score distributions exposed a key distinction between KOLF2.1J mapping representations. While the diploid graph has a mean MAPQ of 54.9, the diploid assembly exhibited a substantially lower mean MAPQ of 6.9, with a significant fraction of reads scoring between 1-10 (Figure 4D). This reflects the effect of mapping to a diploid assembly without merging equivalent sequences, due to ambiguous placements at homozygous regions where reads can map almost equivalently to both haplotypes.

Mapping experiments using long-read sequencing data confirmed these trends (Supplementary Figure 1, Supplementary Table 4). Perhaps surprisingly, given the much longer read lengths involved, we also find that mapping long reads to the diploid assembly rather than the haplotype merged diploid KOLF2.1J graph still results in high levels of mapping ambiguity. Confirming this observation, we see the same mapping quality distribution mapping reads with minimap2. Overall, these results demonstrate that personalized diploid assemblies and genome graphs meaningfully reduce mapping differences relative to traditional linear references and (more marginally) population pangenome references, with direct implications for more reliable variant calling and other secondary applications. Although diploid assemblies and graphs show similar mapping performance, care must be taken when interpreting the MAPQ scores of tools designed for haploid references.

### Cell type-specific, allele-specific, and cell type and haplotype interactive DMRs

DNA methylation is associated with aging (Tharakan et al. 2023; Unnikrishnan et al. 2019)(Bell et al. 2019; Horvath 2013), cell-type identity (Loyfer et al. 2023), and parent-of-origin imprinting (Cuellar Partida et al. 2018)(Court et al. 2014; Akbari et al. 2023). However, the regulatory landscape of DNA methylation at the intersection of cell-type specificity and haplotype resolution remains poorly understood. To address this, we analyzed methylation patterns across five distinct cell types, including four differentiated cell types and iPSCs. Using ONT long-read RNA sequencing data aligned to a personalized diploid reference genome, we identified differentially methylated regions (DMRs) to resolve allele and cell type specific methylation patterns.

We identified 215,689 cell type specific DMRs (cDMRs), with microglial cells exhibiting the highest number among all tested cell types (Figure 5A). As expected, cDMRs at loci corresponding to known cell type marker genes were hypomethylated across both haplotypes, indicating that these genes are likely expressed from both haplotypes (Figure 5B). For example, promoters of *GFAP* (glial fibrillary acidic protein) were specifically hypomethylated in astrocytes, *TREM2* in microglia, and promoters of *POU5F1* and *NANOG* in iPSCs.

**Figure 5.**
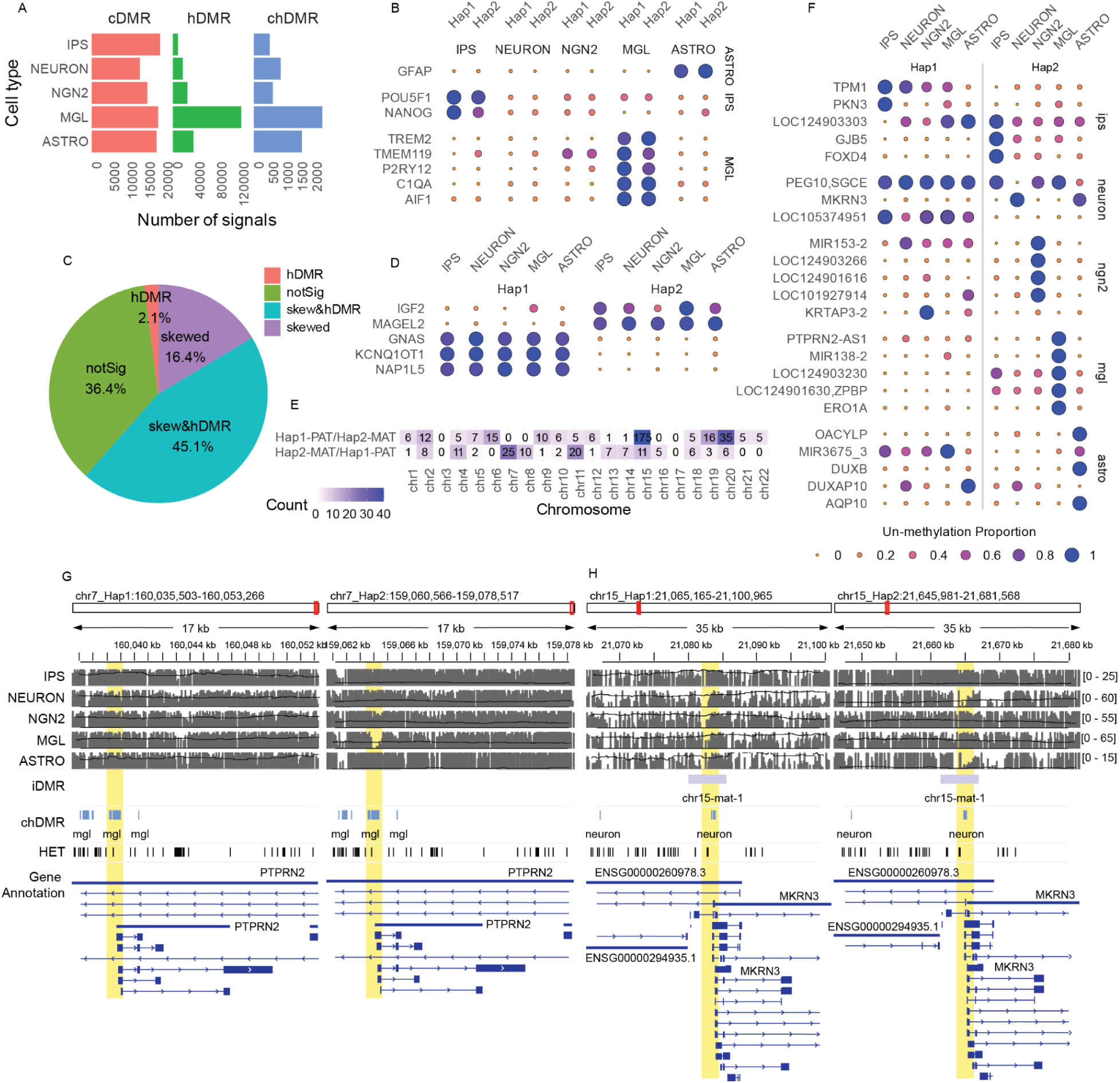
Allele- and cell type–specific DNA methylation landscapes across KOLF2.1J cell types. (A) Number of differentially methylated regions (DMRs) identified for haplotype-specific (hDMRs), cell type specific (cDMRs), and cell type / haplotype interaction effects (chDMRs). (B, D, E) Proportion of unmethylated DMRs overlapping promoter regions (defined as 1,500 bp upstream to 100 bp downstream of transcription start sites) across all cell types. (B) Cell type marker genes. (D) Known imprinted genes. (E) DMRs with haplotype–cell type interaction effects. (C) Proportion of iDMRs intersecting with hDMRs, classified as skewed or not significant. (F) Number of iDMRs exhibiting skewed methylation toward one haplotype, inferred from methylation levels. (F, H) Representative examples of chDMRs displaying allele- and cell type specific hypomethylation patterns. (G) MGL-Hap2-specific chDMR at the promoter of *PTPRN2*. (H) Neuron-Hap2-specific chDMR at the promoter of *MKRN3*. NEURON= Cortical neurons, NGN2= NGN2-induced neurons, MGL=Microglia, ASTRO=Astrocyte-UCL

To assess whether the haplotype-specific DMRs (hDMRs) overlapped with previously characterized imprinted DMRs (iDMRs) (Akbari et al. 2023), we classified regions as “skewed” if the difference in methylation proportion at the region between the two haplotypes exceeded 0.2, “skewed & hDMR” if the region intersected with our *de novo* identified hDMRs, and “not significant” if neither criterion applied (Figure 5C and see Methods). Among the iDMRs, we found that the hDMRs exhibited methylation patterns consistent with known imprinted genes and their respective haplotypes. *KCNQ1OT1*, a well-characterized imprinted gene on chromosome 11, is known to be hypomethylated on the paternal allele (Jima et al. 2022). In the KOLF2.1J genome, *KCNQ1OT1* was likewise hypomethylated in hap1 paternally inherited haplotype, as expected (Figure 5E).

Interestingly, we also identified interaction effects between cell type and haplotype, referred to hereafter as cell type and haplotype-specific DMRs (chDMRs), in which differential methylation between haplotypes was observed only in a specific cell type (Figure 5F). Among the cell types analyzed, microglia exhibited the highest number of chDMRs, followed by astrocytes (Figure 5A). One illustrative example is a chDMR located in the promoter region of *PTPRN2*, which was significantly hypomethylated in the hap2 predicted as paternal of microglial cells, but not in any other haplotype across all cell types (Figure 5G).

We also examined *MKRN3*, a known imprinted gene associated with central precocious puberty (Palumbo et al. 2022; Valadares et al. 2019; Abreu et al. 2013). *MKRN3* acts as a repressor of gonadotropin-releasing hormone (GnRH) release from the hypothalamus, thereby preventing premature activation of the reproductive axis (Abreu et al. 2020, 2015), and it is normally expressed only from the paternally inherited allele (Jima et al. 2022). Notably, we found that the promoter region of *MKRN3* exhibited cell type specific imprinting, with hypomethylation observed exclusively in the neuron paternal hap2 (Figure 5F,H). This suggests that the regulation of MKRN3 may be neuron-specific and implicates neuronal cells as a likely context in which imprinting defects contribute to disease manifestation.

## Discussion

In this study, we generated a complete, high-quality, haplotype-resolved genome assembly for the widely used iPSC line KOLF2.1J and demonstrated its utility across multiple data types: genomic, transcriptomic, and epigenomic. Our results provide compelling evidence that personalized assemblies of commonly used cell lines improve the accuracy of read mapping, variant detection, and epigenomic profiling compared to traditional reliance on generic human reference genomes such as GRCh38. This work represents an important step toward precision iPSC research by ensuring that widely shared cellular models are studied against their own genomic backgrounds.

After polishing, our assembly reached a quality score of Q67.4. This is similar to quality scores reported in previous “gold standard” custom assemblies such as the recent HG002 release (Q68.9)(Hansen et al. 2025). Most of the remaining gaps (99.6%) are attributed by the rDNA arrays, which are the most difficult region to resolve in genome assemblies due to their high repetitiveness and similarity across acrocentric chromosomes. In KOLF2.1Jv1.1, each unresolved rDNA array was placed with a 1Mb gap leading to 10Mb of total missingness.

Genomic mapping experiments revealed that the KOLF2.1J diploid assembly and genome graph consistently reduce mapping differences relative to both traditional linear references (GRCh38, T2T-CHM13) and (to a lesser extent) pangenome references across both short- and long-read datasets. These improvements were most evident in the proportion of perfectly mapped reads and reduced error rates. However, our analysis also highlights the limitations of mapping both short and long reads to diploid assemblies alone, where ambiguity between equivalent haplotypes can reduce the reported mapping quality. In contrast, graph representations of the KOLF2.1J genome mitigated these issues, enabling more confident read placement. Together, these findings emphasize that custom genomes and their graphs represent a powerful tool for downstream analyses, from variant calling to transcript quantification.

Long-read DNA sequencing data also allowed us to catalog around 25,000 high confidence structural variants in KOLF2.1J, including more than 180 that directly affected likely protein coding sequences, which is not a common practice in iPSC line research. Integration with long-read RNA sequencing confirmed that several of these events aligned with transcriptional and gene expression differences, though their functional significance remains unclear. While none of these coding structural variants overlapped a known ADRD gene, the goal of the INDI initiative with KOLF2.1J, this catalog can be of use for future work as the field evolves and more experiments are performed with this line.

Our gene annotation of the KOLF2.1J assembly yielded gene counts on par with CHM13 when using the same pipeline and source gene set, consistent with what is expected from a complete human genome assembly. All single-copy genes from GENCODE were successfully lifted over, with differences between KOLF2.1J and CHM13 driven entirely by copy number variable gene families, which is consistent with known patterns from projects like HPRC (Liao et al. 2023). The annotation also confirms two previously reported CNVs in coding regions, particularly important for studies targeting genes near these loci. Integration of long-read transcriptomic data from five KOLF2.1J-derived cell types identified ∼1,900 novel protein-coding isoforms, including in neurodevelopmentally relevant genes, suggesting that the standard GENCODE catalog does not fully represent the transcript diversity present in this line.

We used ONT methylation data to generate a high resolution map of DNA methylation that resolved both cell type-specific and haplotype specific regulatory signatures. As expected, we observed strong enrichment of hypomethylated regions near canonical cell marker genes, such as *GFAP* in astrocytes and *TREM2* in microglia. Importantly, we also identified thousands of haplotype-specific differentially methylated regions (hDMRs), many of which overlapped known imprinted loci such as *KCNQ1OT1* and *MKRN3*. Our analysis further revealed interaction effects between haplotype and cell type (chDMRs), demonstrating that allele-specific regulation can manifest in different ways.

Collectively, our results argue for the routine generation of personalized genome assemblies for commonly used iPSC lines. These resources not only alleviate reference bias but also serve as a stable foundation for integrating multi-omic data across research groups. To aid in this, we have published the assembly, annotation, and all accompanying genomic, epigenetic, and transcriptomic data in a publicly available browser (https://genome.ucsc.edu/cgi-bin/hgGateway?hgHub_do_redirect=on&hgHubConnect.remakeTrackHub=on&hgHub_do_firstDb=1&hubUrl=https://research.nhgri.nih.gov/CustomTracks/T2T_hubs/T2T_test/KOLF2.1Jv1.1/hub.txt) within the UCSC platform (Casper et al. 2026), in line with other personalized assembly releases like CHM13 and HG002. This will allow any researcher working with the KOLF2.1J to have access to a high quality dataset for KOLF2.1J across multiple cell states and to easily incorporate the assembly and annotation into their own work. Just as the adoption of T2T-CHM13 and pangenome references has reshaped genomic research, we envision that personalized assemblies will become a cornerstone of iPSC disease modeling.

Despite these advances, several limitations remain. First, our genome assembly and annotation provide only a snapshot of KOLF2.1J at a single point in time. iPSCs are known to accumulate genomic changes during culture which may result in line to line variability even within the same original donor (Yoshihara et al. 2017; Liu et al. 2014). Additionally, the mitochondrial genome has not been assessed in this study, where bottle necks occur and will change over time. As such, the KOLF2.1J reference we provide should be seen as a baseline against which future changes can be assessed. Second, while we tried to build the most complete assembly possible, there is still some missingness in hard to resolve areas due to technology limitations. Lastly, while we profiled several neuronal and glial derivatives, many cell types remain unexplored, and the regulatory annotation of KOLF2.1J will continue to improve as more differentiation protocols and omics datasets are applied.

In conclusion, our work establishes the value of personalized genome assemblies for widely used iPSC models. By demonstrating improvements across read mapping, structural variant discovery, and allele-resolved epigenomic profiling, we provide a framework for future efforts to integrate high quality assemblies into iPSC based research. We propose that such personalized reference assemblies should become a routine component of community iPSC projects, analogous to the role of the Genome Reference Consortium genomes, thus advancing precision in disease modeling.

## Methods

### Cell line selection and culture protocols

The iNDI reference parental iPSC KOLF2.1J line was used and obtained from the Jackson Laboratory (Cat# JIPSC001000). Cell pellets were generated and stored at -80C and used for both conventional and high molecular weight DNA and RNA isolations. Previously generated short read Illumina WGS and RNAseq was obtained from Pantazis et al (Pantazis et al. 2022). KOLF2.1J was previously generated and reported to be derived from material obtained under informed consent and appropriate ethical approvals. Y-chromosome data were excluded from this study in accordance with donor consent and data protection requirements. For all cell differentiation protocols please see the supplementary methods.

### iPSC and differentiated cell culture processing

For DNA and RNA extraction and sequencing protocols please see the supplementary methods.

### Sequencing data processing

#### Data processing Oxford Nanopore sequencing data

All the sequencing runs were basecalled on NIH’s HPC (Biowulf) using Oxford Nanopore’s Guppy/6.1.2 in super accuracy mode with the *dna_r9.4.1_450bps_modbases_5mc_cg_sup_prom.cfg* configuration file. Additionally, our N50 30kb whole genome and ultra-long sequencing runs were basecalled with the *–bam_out* option to preserve methylation tags generated by Guppy.

The unmapped bams with methylation tags for the DNA runs were then converted to fastqs using Samtools/1.17 (Danecek et al. 2021) (*samtools fastq -TMm, Ml*) and initially mapped to hg38 using Minimap2/2.26 (Li 2018) with ONT defaults. For information on mapping to the personalized assembly please see custom mapping section below.

To call variants, we used PEPPER-Margin-DeepVariant/0.8 (Shafin et al. 2021) to call small variants (<50bp) and phase our variant calls and alignment. We then used our phased alignment, to produce haplotype-specific methylation calls using Modbamtools/0.4.8 (Razaghi et al. 2022) and Nanopore’s modbam2bed (GitHub, n.d.-a).

After basecalling, our RNA files were mapped to hg38 and our KOLF assembly using Minimap2/2.26 (Li 2018) with splice aware parameters (*-ax splice -k 14 -uf*). Transcripts were then called using Stringtie2 with the following parameters: stringtie --rf -G ${annotation} -L -v -p 10 -A ${prefix}_gene_abundance.tab -o ${prefix}.gtf ${input}.

#### Data processing PacBio sequencing data

Basecalled files were provided to us by Psomagen following standard PacBio guidelines.

### Custom genome generation and subsequent analyses

#### Genome assembly and polishing

##### Assembly

An initial assembly (v0.6) was produced with Verkko (v1.4) using the HiFi and ONT Ultra-long reads. Subsequently, the assembly graph was manually curated over several rounds for better hi-c integration and traversing gaps. In the first round, 32 paths have been manually curated using ONT alignments to the graph Verkko couldn’t automatically resolve as described in Yoo et al (Yoo et al. 2025). In brief, ONT reads were re-aligned to the final graph using GraphAligner v1.0.17. Depth of coverage and node length were considered to remove artifactual edges and resolve simple nonlinear structures. For more details, refer to Supplementary Information “II. Assembly” part of Yoo et al.

##### Chromosome assignment

Each sequence was mapped to CHM13v2.0 with mashmap3 using options -r chm13v2.0.fa -q assembly.fasta -s 1000000 --pi 90 -t 24 for chromosome assignment. Sequences over 1Mbp were considered for chromosome assignments. A sequence was assigned to a chromosome if more than 50% of the sequence(s) mapped to one chromosome. Most chromosomes came out as T2T one contiguous sequence (contig) with no gaps, except for the acrocentric chromosomes 13, 14, 15, 21, 22, chromosome 10 (both haplotypes), and one haplotype of chromosomes 1, 7, 17 and 19. Sequence orientation was evaluated compared to the assigned chromosome and reverse complemented for those needed with samtools faidx -r -l seq_to_rev.list. Haplotypes 1 and 2 were temporarily assigned based on the sequence names given by verkko at this stage, pending polishing and patching improvements for the final haplotype assignments.

Using mashmap results, one sequence (unassigned-0000682) was selected and assigned to the mitochondria based on mapped % identity and sequence length.

#### Polishing

##### Single Nucleotide Variant Correction

A variant calling based SNV correction was applied similarly as described in McCartney et al., 2023 (Mc Cartney et al. 2022) and Yoo et al., 2025 (Yoo et al. 2025). In brief, a diploid and haploid version of the KOLF assembly was prepared. The diploid version (analysis-dip for brevity) contains both haplotype autosomes and sex chromosomes and the Mitochondrial sequence. The haploid version (hap for brevity) hap1 contains one haplotype of chromosomes 1-22 + X, while hap2 contains the other haplotype 2 of chromosomes 1-22 + Y, respectively. PacBio HiFi and ONT UL reads were mapped with winnowmap2 v2.03 (Jain et al. 2022) using -I 8G --MD -W repetitive_k15.txt -ax $map -t$cpus $ref $reads. For $map, map-pb was used for HiFi and map-ont for ONT reads. Resulting sam file was sorted with samtools v1.21 sort -@$cpus -m2G -T $tmp/$out.tmp -O bam -o $tmp/$out.sort.bam $tmp/$out.sam. Illumina short-reads were mapped with bwa v0.7.17 [REF: PMID 19451168] mem -t $cpus $ref $r1 $r2, and subsequently processed with samtools fixmate -m -@$cpu $tmp/$out.sam $tmp/$out.fix.bam, samtools sort -@$cpu -O bam -o $tmp/$out.bam -T $tmp/$out.tmp $tmp/$out.fix.bam, and samtools markdup -r -@$cpu $tmp/$out.bam $tmp/$out.dedup.bam. After removing secondary alignments with samtools view -@$cpu -F0x100 -hb --write-index, the primary alignments were used to generate a hybrid bam file with the HiFi mappings described above.

Variant calling was performed with DeepVariant v1.5 (Poplin et al. 2018) in PACBIO, HYBRID, and ONT R9 mode accordingly in default mode except for the dip alignments, whereas all reads including MQ0 were used in step 1 of DeepVariant.

A hybrid 31-mer database was generated from the Illumina and PacBio HiFi reads for further filtering SNV correction candidates with Merqury v1.4.1 (Rhie et al. 2020) and Merfin v1.1 (Formenti et al. 2022). In brief, 31-mers were collected with Meryl v1.4.1 using commands meryl count -k31, and merged with meryl union-sum [ greater-than 1 hifi.k31.meryl ] [ greater-than 1 illumina.k31.meryl ]. This considers only kmers seen at least twice in each read set to determine the correction is supported by the reads. The final variants with PASS assigned by DeepVariant were considered and processed with Merfin. The full code can be found on our Github.

##### Structural Variant Correction

A second round of assembly (v0.7) was performed using the same reads using Verkko v2.0, as many improvements have been applied in the newer version of Verkko for better integration of the Hi-C scaffolding and resolving complex repeats. The new assembly was aligned to the initial graph-curated assembly with minimap2 v2.24 -x asm5 option (Li 2018) to find the corresponding distal bits and proximal bits from the rDNA. Candidate sequences to replace regions with possible remaining mis-assemblies or closable gaps were identified.

If the new sequence contained gaps, the sequence was broken at gaps to prevent mis-alignments. Once the pairing region was identified in a chr_hap pair, for the prior and the newer assembly, wfmash --no-split -ad -s100000 -p95 --one-to-one was used to get the best matching alignments to v0.6 for each chr_hap of interest. Each aligned bam file was then evaluated with IGV v2.18.2, and the hybrid 31-mers were used to flag excessive error k-mers in the patch sequence. The patch region was selected based on best concordance (∼1Mb around the region of interest) to ensure the mapping was reliable, and by excluding regions with error kmers to minimize the introduction of additional errors. Patch sequences were then generated in a VCF format, to contain the original sequence in REF field, and the replacement sequence in ALT field (KOLF2.1Jv0.6.patch.vcf.gz).

##### Final consensus

Larger regions where manual patches were identified either from a newer version of assembly or through re-walking the assembly graph were recorded in patch.bed and removed from the final SNV polishing candidates. Patches were recorded as SV correction VCF and concatenated along with the rest of the SNV candidates. The final consensus was generated with bcftools consensus as below and re-evaluated.

bedtools subtract -a v0.6_chrM.bed -b patch.bed > v0.6_include_for_polishing.bed

bcftools view -R v0.6_include_for_polishing.bed --no-version --threads 24 -Oz v0.6_snv_candidates.merfin-loose.vcf.gz > v0.6_snv_correction.vcf.gz

bcftools index v0.6_snv_correction.vcf.gz

bcftools concat --no-version --threads 24 -a -Oz v0.6_snv_correction.vcf.gz KOLF2.1Jv0.6.patch.vcf.gz > v0.6_snv_sv_correction.vcf.gz

bcftools index v0.6_snv_sv_correction.vcf.gz

bcftools consensus -c v0.6_to_$new_ver.chain -HA -f v0.6_chrM.fa v0.6_snv_sv_correction.vcf.gz > $new_ver.fa

At last, the haplotypes were re-assigned so the more contiguous (gapless), higher quality haplotype becomes the primary assembly (hap1). The Y chromosome was excluded from release to abide by the KOLF cell line consent.

##### Validation

Base level accuracy before and after polishing was measured using Illumina 31-mers and hybrid 31-mers with Merqury. Structural validation was performed using long-read alignments to the v1.1 assembly. Reads were mapped with Winnowmap v2.03 using k=15 for downweighting repetitive kmers and options -x map-pb for hifi and -x map-ont for ONT reads as recommended (Mc Cartney et al. 2022). After excluding secondary alignments, structural discrepancies with reads were collected with the T2T-Polish coverage-based evaluation pipeline (Mc Cartney et al. 2022). To avoid sequencing bias, regions called as misassembly in both HiFi and ONT were considered and reported as the final issues. Using the pipeline, regions with excessive clipped read alignments (> 10% in HiFi and > 15% in ONT), low (< mean / 4) or high (> mean x 2.5) coverage were collected for each platform. The two “issues” files were intersected with `bedtools intersect -u -a $hifi_issues -b $ont_issues`, and the gap regions were added after distinguishing the rDNA gaps and others. The final bed file is present in Supplementary Table 1.

#### Structural variants of the diploid KOLFv1.1 assembly

To catalog structural variants we ran hapdiff on the diploid polished assembly (https://github.com/KolmogorovLab/hapdiff). We ran hapdiff with standard parameters and then filtered the call by a minimum length of 50bp and removed any calls in centromeres or satellite regions per GIAB stratification files (https://ftp-trace.ncbi.nlm.nih.gov/ReferenceSamples/giab/release/references/GRCh38/resources/) using bcftools (Danecek et al. 2021). The SVs were then annotated using both AnnotSV (Geoffroy et al. 2018) and Annovar (Wang et al. 2010) against GrCh38 using standard parameters.

#### Custom Genome annotation

Genome annotation was performed using the Comparative Annotation Toolkit (CAT) (improving upon (Fiddes et al. 2018)). First, whole-genome alignments between the two KOLF2.1J haplotypes, GRCh38, and CHM13 genomes were generated using Cactus (Armstrong et al. 2020). CAT then used the whole-genome alignments to project the GENCODE v44 annotation set from GRCh38 to the KOLF2.1J haplotypes. CAT was run with transMap, AUGUSTUS, Liftoff, AUGUSTUS-PB, and miniprot modes. transMap lifts over gene annotations from the reference onto all the genomes in the cactus alignment. Liftoff lifts over gene annotations from a reference onto a minimap2 alignment between the reference and target genome. The miniprot mode uses protein homology information to improve gene annotations. CAT was given ONT RNA data to provide extrinsic hints for ab initio prediction of coding isoforms. CAT then combined these ab initio prediction sets with the various gene projection sets to produce the final gene sets used in this project. Novel isoforms were identified by comparing transcript models assembled from the long-read data against the projected GENCODE set. Transcripts that showed novel splice junctions, exon skipping events, or previously unannotated exon combinations not present in the reference annotation were flagged as candidate novel isoforms.

#### Mapping of sequencing reads to a custom genome build

To evaluate the performance of custom genome mapping versus widely used reference genomes, we conducted comparative mapping analyses using KOLF2.1J paired-end Illumina short-read data (866 million reads) and PacBio long-read HiFi data (9.9 million reads). Our comparison encompassed six mapping targets, including two linear references, CHM13 (Nurk et al. 2022) and GRCh38 (Schneider et al. 2017), the HPRC v2.0 pangenome, an HPRC v2.0 personalized pangenome, the KOLF2.1J diploid assembly (in which both maternal and paternal haplotypes are included as separate sequences), and a custom KOLF2.1J diploid graph (in which the maternal and paternal haplotypes are merged where they are equivalent).

The KOLF2.1J diploid graph was constructed by integrating KOLF2.1J paternal and maternal haplotypes with the CHM13 using Minigraph-Cactus (Hickey et al. 2023), generating a graph-based representation that captures KOLF2.1J-specific variation. By including CHM13 in the graph we allow conversion to standard reference coordinates (Figure 4A). The personalized pangenome was generated using haplotype sampling (Sirén et al. 2024) from the HPRC v2.0 pangenome, which selected optimal pangenome haplotypes based on variation patterns present in the KOLF2.1J sequencing reads (Figure 4A).

All mappings to the graph as well as linear references and the diploid assembly were performed using vg giraffe, the only mapper to our knowledge capable of mapping both long and short reads to all these references (Sirén et al. 2021), refer to supplementary methods for details. Subsequently, we analyzed mapping statistics across all reference types, including mapping rates and alignment quality scores, to assess the relative performance of these different mapping substrates.

### Processing and analysing methylation data of KOLF2.1J cell types

#### Calling methylation

ONT reads were aligned to the diploid reference using winnowmap v2.03 with the -y -Y parameters to keep the methylation tags (Jain et al. 2020). High-frequency *k*-mers in the diploid reference (KOLF2.1J), downweighted by winnowmap during alignment seeding, were identified with meryl v1.4.1 (Rhie et al. 2020). The resulting alignment files were merged, and only primary alignments were retained using samtools v1.21. Methylation calling and coverage count for each position was performed using modkit (v0.5.0) in pileup mode using default settings (GitHub, n.d.-c).

#### Generation of chain files between haplotypes

We used the nf-LO pipeline v1.8.0 (Nextflow v24.10.3) with minimap2 v2.28 to align source and target assemblies and generate chain files (Talenti and Prendergast 2021). Chain files were split and converted to PAF format using chaintools v0.4 (GitHub, n.d.-b), then cleaned with the rb utility to break large blocks and trim low-quality ends. The refined PAF was converted back into chain format with paf2chain v0.1.0 (Guarracino, n.d.) and inverted chain files were generated using invert.py from the chaintools library. The final outputs were chain files used for coordinate liftover between assemblies. For chains between KOLF haplotypes and CHM13, we used KOLF2.1J as the source and CHM13 as the target for both haplotypes. For chains between KOLF haplotypes, we used the primary haplotype as the source and the alternative haplotype as the target.

#### Differentially methylated regions (DMR) analysis

Methylation counts were lifted over from each haplotype to the CHM13v2.0 coordinates using CrossMap v0.7.0 with the corresponding chain file. This was performed separately for each haplotype across all cell types (Zhao et al. 2014). Only autosomes were included in the analysis. DMRs specific to cell type, allele, and their interaction were identified using a multi-factor linear modeling framework implemented in DSS v2.48.0 (Wu et al. 2013). Methylated and unmethylated read counts at each CpG site, across all haplotypes and cell types, were loaded into a BSseq object. For each cell type, a linear regression model was fitted by labeling the target cell type as the ’case’ and all other cell types as ’control’, enabling the detection of DMRs distinguishing each cell type from the rest.

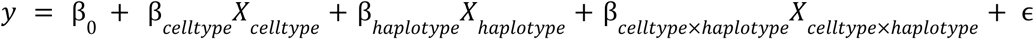

The y-values represent the methylation level at each CpG site, estimated using the DMLfit.multiFactor function. This model was subsequently passed to the DMLtest.multiFactor function to perform a Wald test at each CpG site. Finally, the callDMR function was used to identify DMRs based on the DML test results, using a p-value threshold of 0.05.

To identify DMRs located within promoter regions, transcription start sites (TSSs) were defined as the region spanning 1500 bp upstream to 500 bp downstream of the annotated start site of each transcript, based on CHM13v2.0 gene annotation (https://s3-us-west-2.amazonaws.com/human-pangenomics/T2T/CHM13/assemblies/annotation/chm13v2.0_RefSeq_Liftoff_v5.2.gff3.gz). DMRs overlapping these TSS regions were extracted, and gene names were assigned based on the intersecting transcript annotations.

#### Inferring parent-of-origin haplotypes

We downloaded known iDMR coordinates based on CHM13 from Akbari et al. 2023 (Akbari et al. 2023) and calculated the methylation ratio for each region as the total number of modified bases divided by the total coverage. This was done for both haplotypes lifted to CHM13 coordinates. To quantify methylation bias, we defined the skew of methylation as the minimum methylation ratio divided by the maximum methylation ratio between haplotypes for each iDMR. Regions were classified as “skewed” if the skew value exceeded 0.2, and as “not skewed” otherwise.

We used these skewed iDMRs to infer the parental origin of haplotypes for each chromosome and contig. Specifically, we compared two scenarios—(1) hap1 haplotype as maternal and hap2 haplotype as paternal, and (2) the reverse assignment—and determined the parent of origin based on which scenario yielded more skewed iDMRs consistent with the known haplotype-specific methylation patterns.

## Author contributions

Conceptualization: PAJ, BP, AP, CB

Experiments were performed by: PAJ, EL, AB, CA, JZ, PWC, DP, LM, FG

Data was provided by: PAJ, EL, AB, CA, JZ, WCS, KB, VF, SW, MW

Data was analyzed by: PAJ, AR, JK, PB, SN, DA, SK, NFH, GH, BW

Writing (original draft): PAJ, AR, JK, PB, SN

Writing (review and editing): PAJ, AR, JK, PB, DA, SK, EL, AB, NFH, CA, JZ, PWC, DP, GH, BW, LM, WCS, XR, RG, KD, CBP, FG, MAN, KB, VF, SW, MW, MR, ABS, MRC, MJ, BP, AP

## Code Availability

All scripts and code for this project, including assembly polishing, assembly annotation, SV calling, mapping, and methylation can be found at: https://github.com/pilaralv/KOLF2.1J_ASSEMBLY

## Data Availability

T2T-browser : https://github.com/marbl/T2T-Browser?tab=readme-ov-file

KOLF browser : https://genome.ucsc.edu/cgi-bin/hgGateway?hgHub_do_redirect=on&hgHubConnect.remakeTrackHub=on&hgHub_do_firstDb=1&hubUrl=https://research.nhgri.nih.gov/CustomTracks/T2T_hubs/T2T_test/KOLF2.1Jv1.1/hub.txt

All resources and data generated for this project can be found at the following links : https://github.com/marbl/KOLF2.1J-Diploid-Assembly

## Funding agencies

This work was supported in part by the Intramural Research Program of the National Institutes of Health including: the Center for Alzheimer’s and Related Dementias, within the Intramural Research Program of the National Institute on Aging and the National Institute of Neurological Disorders and Stroke (1ZIAAG000538-03, ZIAAG000542-01 and 1ZIAAG000543-01), and the National Human Genome Research Institute. The contributions of NIH authors are considered Works of the United States Government. The findings and conclusions presented in this paper are those of the authors and do not necessarily reflect the views of the NIH or the U.S. Department of Health and Human Services. This research was funded in part by Aligning Science Across Parkinson’s MJFF-026403 and MJFF-024547 through the Michael J. Fox Foundation for Parkinson’s Research (MJFF). This work utilized the computational resources of the NIH HPC Biowulf cluster (http://hpc.nih.gov). S.W. and C.A. were supported by the National Institute for Health and Care Research University College London Hospitals Biomedical Research Centre.

## References

1. “2022 Alzheimer’s Disease Facts and Figures.” 2022. Alzheimer’s & Dementia: The Journal of the Alzheimer’s Association 18 (4): 700–789.

2. Abreu, Ana Paula, Andrew Dauber, Delanie B. Macedo, et al. 2013. “Central Precocious Puberty Caused by Mutations in the Imprinted Gene MKRN3.” The New England Journal of Medicine 368 (26): 2467–2475.

3. Abreu, Ana Paula, Delanie B. Macedo, Vinicius N. Brito, Ursula B. Kaiser, and Ana Claudia Latronico. 2015. “A New Pathway in the Control of the Initiation of Puberty: The MKRN3 Gene.” Journal of Molecular Endocrinology 54 (3): R131–9.

4. Abreu, Ana Paula, Carlos A. Toro, Yong Bhum Song, et al. 2020. “MKRN3 Inhibits the Reproductive Axis through Actions in Kisspeptin-Expressing Neurons.” The Journal of Clinical Investigation 130 (8): 4486–4500.

5. Abud, Edsel M., Ricardo N. Ramirez, Eric S. Martinez, et al. 2017. “iPSC-Derived Human Microglia-like Cells to Study Neurological Diseases.” Neuron 94 (2): 278–293.e9.

6. Aganezov, Sergey, Stephanie M. Yan, Daniela C. Soto, et al. 2022. “A Complete Reference Genome Improves Analysis of Human Genetic Variation.” *Science*, ahead of print, April 1. 10.1126/science.abl3533.

7. Akbari, Vahid, Vincent C. T. Hanlon, Kieran O’Neill, et al. 2023. “Parent-of-Origin Detection and Chromosome-Scale Haplotyping Using Long-Read DNA Methylation Sequencing and Strand-Seq.” Cell Genomics 3 (1): 100233.

8. Antipov, Dmitry, Mikko Rautiainen, Sergey Nurk, et al. 2025. “Verkko2 Integrates Proximity-Ligation Data with Long-Read De Bruijn Graphs for Efficient Telomere-to-Telomere Genome Assembly, Phasing, and Scaffolding.” Genome Research 35 (7): 1583–1594.

9. Armstrong, Joel, Glenn Hickey, Mark Diekhans, et al. 2020. “Progressive Cactus Is a Multiple-Genome Aligner for the Thousand-Genome Era.” Nature 587 (7833): 246–251.

10. Bell, Christopher G., Robert Lowe, Peter D. Adams, et al. 2019. “DNA Methylation Aging Clocks: Challenges and Recommendations.” Genome Biology 20 (1): 249.

11. Casper, Jonathan, Matthew L. Speir, Brian J. Raney, et al. 2026. “The UCSC Genome Browser Database: 2026 Update.” Nucleic Acids Research 54 (D1): D1331–D1335.

12. Cheng, Haoyu, Mobin Asri, Julian Lucas, Sergey Koren, and Heng Li. 2024. “Scalable Telomere-to-Telomere Assembly for Diploid and Polyploid Genomes with Double Graph.” Nature Methods 21 (6): 967–970.

13. Cheng, Haoyu, Gregory T. Concepcion, Xiaowen Feng, Haowen Zhang, and Heng Li. 2021. “Haplotype-Resolved de Novo Assembly Using Phased Assembly Graphs with Hifiasm.” Nature Methods 18 (2): 170–175.

14. Court, Franck, Chiharu Tayama, Valeria Romanelli, et al. 2014. “Genome-Wide Parent-of-Origin DNA Methylation Analysis Reveals the Intricacies of Human Imprinting and Suggests a Germline Methylation-Independent Mechanism of Establishment.” Genome Research 24 (4): 554–569.

15. Cuellar Partida, Gabriel, Charles Laurin, Susan M. Ring, et al. 2018. “Genome-Wide Survey of Parent-of-Origin Effects on DNA Methylation Identifies Candidate Imprinted Loci in Humans.” Human Molecular Genetics 27 (16): 2927–2939.

16. Danecek, Petr, James K. Bonfield, Jennifer Liddle, et al. 2021. “Twelve Years of SAMtools and BCFtools.” GigaScience 10 (2): giab008.

17. Degner, Jacob F., John C. Marioni, Athma A. Pai, et al. 2009. “Effect of Read-Mapping Biases on Detecting Allele-Specific Expression from RNA-Sequencing Data.” Bioinformatics (Oxford, England) 25 (24): 3207–3212.

18. Fiddes, Ian T., Joel Armstrong, Mark Diekhans, et al. 2018. “Comparative Annotation Toolkit (CAT)—simultaneous Clade and Personal Genome Annotation.” Genome Research 28 (7): 1029–1038.

19. Formenti, Giulio, Arang Rhie, Brian P. Walenz, et al. 2022. “Merfin: Improved Variant Filtering, Assembly Evaluation and Polishing via K-Mer Validation.” Nature Methods 19 (6): 696–704.

20. Garrison, Erik, Jouni Sirén, Adam M. Novak, et al. 2018. “Variation Graph Toolkit Improves Read Mapping by Representing Genetic Variation in the Reference.” Nature Biotechnology 36 (9): 875–879.

21. Geoffroy, Véronique, Yvan Herenger, Arnaud Kress, et al. 2018. “AnnotSV: An Integrated Tool for Structural Variations Annotation.” *Bioinformatics (Oxford*, England) 34 (20): 3572–3574.

22. GitHub. n.d.-a. “GitHub - epi2me-labs/modbam2bed.” Accessed May 29, 2023. https://github.com/epi2me-labs/modbam2bed.

23. GitHub. n.d.-b. “GitHub - Lightninglabs/chantools: A Loose Collection of Tools All Somehow Related to Lnd and Lightning Network Channels.” Accessed October 6, 2025. https://github.com/lightninglabs/chantools.

24. GitHub. n.d.-c. “GitHub - Nanoporetech/modkit: A Bioinformatics Tool for Working with Modified Bases.” Accessed October 6, 2025. https://github.com/nanoporetech/modkit.

25. Guarracino, Andrea. n.d. AndreaGuarracino/paf2chain: v0.1.0. 10.5281/zenodo.8108447.

26. Hansen, Nancy F., Nathan Dwarshuis, Hyun Joo Ji, et al. 2025. “A Complete Diploid Human Genome Benchmark for Personalized Genomics.” In bioRxiv. September 21. 10.1101/2025.09.21.677443.

27. Hickey, Glenn, Jean Monlong, Jana Ebler, et al. 2023. “Pangenome Graph Construction from Genome Alignments with Minigraph-Cactus.” Nature Biotechnology 42 (4): 663–673.

28. Horvath, Steve. 2013. “DNA Methylation Age of Human Tissues and Cell Types.” Genome Biology 14 (10): R115.

29. Jain, Chirag, Arang Rhie, Nancy F. Hansen, Sergey Koren, and Adam M. Phillippy. 2022. “Long-Read Mapping to Repetitive Reference Sequences Using Winnowmap2.” Nature Methods 19 (6): 705–710.

30. Jain, Chirag, Arang Rhie, Haowen Zhang, et al. 2020. “Weighted Minimizer Sampling Improves Long Read Mapping.” Bioinformatics (Oxford, England) 36 (Suppl_1): i111–i118.

31. Jain, Miten, Sergey Koren, Karen H. Miga, et al. 2018. “Nanopore Sequencing and Assembly of a Human Genome with Ultra-Long Reads.” Nature Biotechnology 36 (4): 338–345.

32. Jima, Dereje D., David A. Skaar, Antonio Planchart, et al. 2022. “Genomic Map of Candidate Human Imprint Control Regions: The Imprintome.” Epigenetics 17 (13): 1920–1943.

33. Kolmogorov, Mikhail, Kimberley J. Billingsley, Mira Mastoras, et al. 2023. “Scalable Nanopore Sequencing of Human Genomes Provides a Comprehensive View of Haplotype-Resolved Variation and Methylation.” bioRxiv : The Preprint Server for Biology, ahead of print, April 5. 10.1101/2023.01.12.523790.

34. Liao, Wen-Wei, Mobin Asri, Jana Ebler, et al. 2023. “A Draft Human Pangenome Reference.” Nature 617 (7960): 312–324.

35. Li, Heng. 2018. “Minimap2: Pairwise Alignment for Nucleotide Sequences.” Bioinformatics 34 (18): 3094–3100.

36. Lin, Mao-Jan, Sheila Iyer, Nae-Chyun Chen, and Ben Langmead. 2024. “Measuring, Visualizing, and Diagnosing Reference Bias with Biastools.” Genome Biology 25 (1): 1–28.

37. Liu, Pengfei, Anna Kaplan, Bo Yuan, Jacob H. Hanna, James R. Lupski, and Orly Reiner. 2014. “Passage Number Is a Major Contributor to Genomic Structural Variations in Mouse iPSCs.” Stem Cells (Dayton, Ohio) 32 (10): 2657–2667.

38. Loyfer, Netanel, Judith Magenheim, Ayelet Peretz, et al. 2023. “A DNA Methylation Atlas of Normal Human Cell Types.” Nature 613 (7943): 355–364.

39. Mc Cartney, Ann M., Kishwar Shafin, Michael Alonge, et al. 2022. “Chasing Perfection: Validation and Polishing Strategies for Telomere-to-Telomere Genome Assemblies.” Nature Methods 19 (6): 687–695.

40. Nurk, Sergey, Sergey Koren, Arang Rhie, et al. 2022. “The Complete Sequence of a Human Genome.” Science 376 (6588): 44–53.

41. Palumbo, Stefania, Grazia Cirillo, Francesca Aiello, Alfonso Papparella, Emanuele Miraglia Del Giudice, and Anna Grandone. 2022. “MKRN3 Role in Regulating Pubertal Onset: The State of Art of Functional Studies.” Frontiers in Endocrinology 13 (September): 991322.

42. Pantazis, Caroline B., Andrian Yang, Erika Lara, et al. 2022. “A Reference Human Induced Pluripotent Stem Cell Line for Large-Scale Collaborative Studies.” Cell Stem Cell 29 (12): 1685–1702.e22.

43. Poplin, Ryan, Pi-Chuan Chang, David Alexander, et al. 2018. “A Universal SNP and Small-Indel Variant Caller Using Deep Neural Networks.” Nature Biotechnology 36 (10): 983–987.

44. Ranallo-Benavidez, T. Rhyker, Yue Hao, Emilia Volpe, et al. 2025. “Diploid Genome Assembly of Human Fibroblast Cell Lines Enables Clone Specific Variant Calling, Improved Read Mapping and Accurate Phasing.” In bioRxiv. April 15. 10.1101/2025.04.15.648829.

45. Rautiainen, Mikko, Sergey Nurk, Brian P. Walenz, et al. 2023. “Telomere-to-Telomere Assembly of Diploid Chromosomes with Verkko.” Nature Biotechnology, ahead of print, February 16. 10.1038/s41587-023-01662-6.

46. Razaghi, Roham, Paul W. Hook, Shujun Ou, et al. 2022. “Modbamtools: Analysis of Single-Molecule Epigenetic Data for Long-Range Profiling, Heterogeneity, and Clustering.” In bioRxiv. July 8. 10.1101/2022.07.07.499188.

47. Reitz, Christiane, Margaret A. Pericak-Vance, Tatiana Foroud, and Richard Mayeux. 2023. “A Global View of the Genetic Basis of Alzheimer Disease.” Nature Reviews Neurology 19 (5): 261–277.

48. Rhie, Arang, Brian P. Walenz, Sergey Koren, and Adam M. Phillippy. 2020. “Merqury: Reference-Free Quality, Completeness, and Phasing Assessment for Genome Assemblies.” Genome Biology 21 (1): 245.

49. Rozowsky, Joel, Jiahao Gao, Beatrice Borsari, et al. 2023. “The EN-TEx Resource of Multi-Tissue Personal Epigenomes & Variant-Impact Models.” Cell 186 (7): 1493–1511.e40.

50. Schneider, V. A., T. Graves-Lindsay, K. Howe, et al. 2017. “Evaluation of GRCh38 and de Novo Haploid Genome Assemblies Demonstrates the Enduring Quality of the Reference Assembly.” Genome Research 27 (5). 10.1101/gr.213611.116.

51. Shafin, Kishwar, Trevor Pesout, Pi-Chuan Chang, et al. 2021. “Haplotype-Aware Variant Calling with PEPPER-Margin-DeepVariant Enables High Accuracy in Nanopore Long-Reads.” Nature Methods 18 (11): 1322–1332.

52. Sirén, Jouni, Parsa Eskandar, Matteo Tommaso Ungaro, et al. 2024. “Personalized Pangenome References.” Nature Methods 21 (11): 2017–2023.

53. Sirén, Jouni, Jean Monlong, Xian Chang, et al. 2021. “Pangenomics Enables Genotyping of Known Structural Variants in 5202 Diverse Genomes.” Science, ahead of print, December 17. 10.1126/science.abg8871.

54. Sullivan, Sarah E., and Tracy L. Young-Pearse. 2017. “Induced Pluripotent Stem Cells as a Discovery Tool for Alzheimer’s Disease.” Brain Research 1656 (February): 98–106.

55. Talenti, Andrea, and James Prendergast. 2021. “Nf-LO: A Scalable, Containerized Workflow for Genome-to-Genome Lift Over.” Genome Biology and Evolution 13 (9). 10.1093/gbe/evab183.

56. Taylor, Dylan J., Jordan M. Eizenga, Qiuhui Li, et al. 2024. “Beyond the Human Genome Project: The Age of Complete Human Genome Sequences and Pangenome References.” Annual Review of Genomics and Human Genetics 25 (1): 77–104.

57. Tharakan, Ravi, Ceereena Ubaida-Mohien, Christopher Dunn, et al. 2023. “Whole-Genome Methylation Analysis of Aging Human Tissues Identifies Age-Related Changes in Developmental and Neurological Pathways.” Aging Cell 22 (7): e13847.

58. Unnikrishnan, Archana, Willard M. Freeman, Jordan Jackson, Jonathan D. Wren, Hunter Porter, and Arlan Richardson. 2019. “The Role of DNA Methylation in Epigenetics of Aging.” Pharmacology & Therapeutics 195 (March): 172–185.

59. Valadares, Luciana Pinto, Cinthia Gabriel Meireles, Isabela Porto De Toledo, et al. 2019. “Mutations in Central Precocious Puberty: A Systematic Review and Meta-Analysis.” Journal of the Endocrine Society 3 (5): 979–995.

60. Volpe, Emilia, Alessio Colantoni, Luca Corda, et al. 2025. “The Reference Genome of the Human Diploid Cell Line RPE-1.” Nature Communications 16 (1): 7751.

61. Wang, Chao, Michael E. Ward, Robert Chen, et al. 2017. “Scalable Production of iPSC-Derived Human Neurons to Identify Tau-Lowering Compounds by High-Content Screening.” Stem Cell Reports 9 (4): 1221–1233.

62. Wang, Kai, Mingyao Li, and Hakon Hakonarson. 2010. “ANNOVAR: Functional Annotation of Genetic Variants from High-Throughput Sequencing Data.” Nucleic Acids Research 38 (16): e164.

63. Wenger, A. M., P. Peluso, W. J. Rowell, et al. 2019. “Accurate Circular Consensus Long-Read Sequencing Improves Variant Detection and Assembly of a Human Genome.” Nature Biotechnology 37 (10). 10.1038/s41587-019-0217-9.

64. Wu, Hao, Chi Wang, and Zhijin Wu. 2013. “A New Shrinkage Estimator for Dispersion Improves Differential Expression Detection in RNA-Seq Data.” Biostatistics (Oxford, England) 14 (2): 232–243.

65. Yoo, Dongahn, Arang Rhie, Prajna Hebbar, et al. 2025. “Complete Sequencing of Ape Genomes.” Nature 641 (8062): 401–418.

66. Yoshihara, Masahito, Yoshihide Hayashizaki, and Yasuhiro Murakawa. 2017. “Genomic Instability of iPSCs: Challenges Towards Their Clinical Applications.” Stem Cell Reviews and Reports 13 (1): 7–16.

67. Zhao, Hao, Zhifu Sun, Jing Wang, Haojie Huang, Jean-Pierre Kocher, and Liguo Wang. 2014. “CrossMap: A Versatile Tool for Coordinate Conversion between Genome Assemblies.” Bioinformatics (Oxford, England) 30 (7): 1006–1007.

