## Supplementary Figure 1 for "The complete genome of the KOLF2.1J reference iPSC line"

### Slide 1
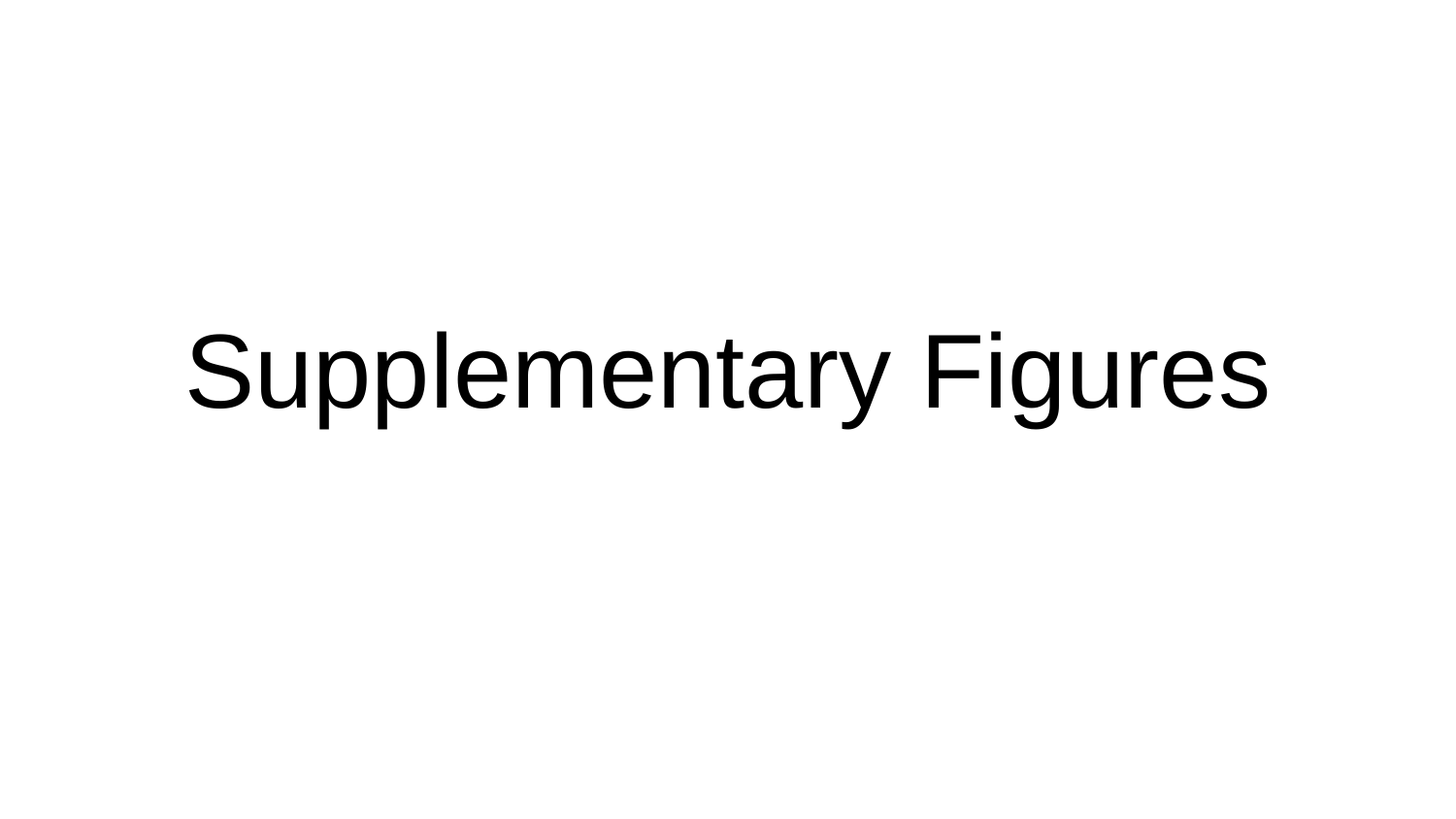

#### Supplementary Figures

### Slide 2
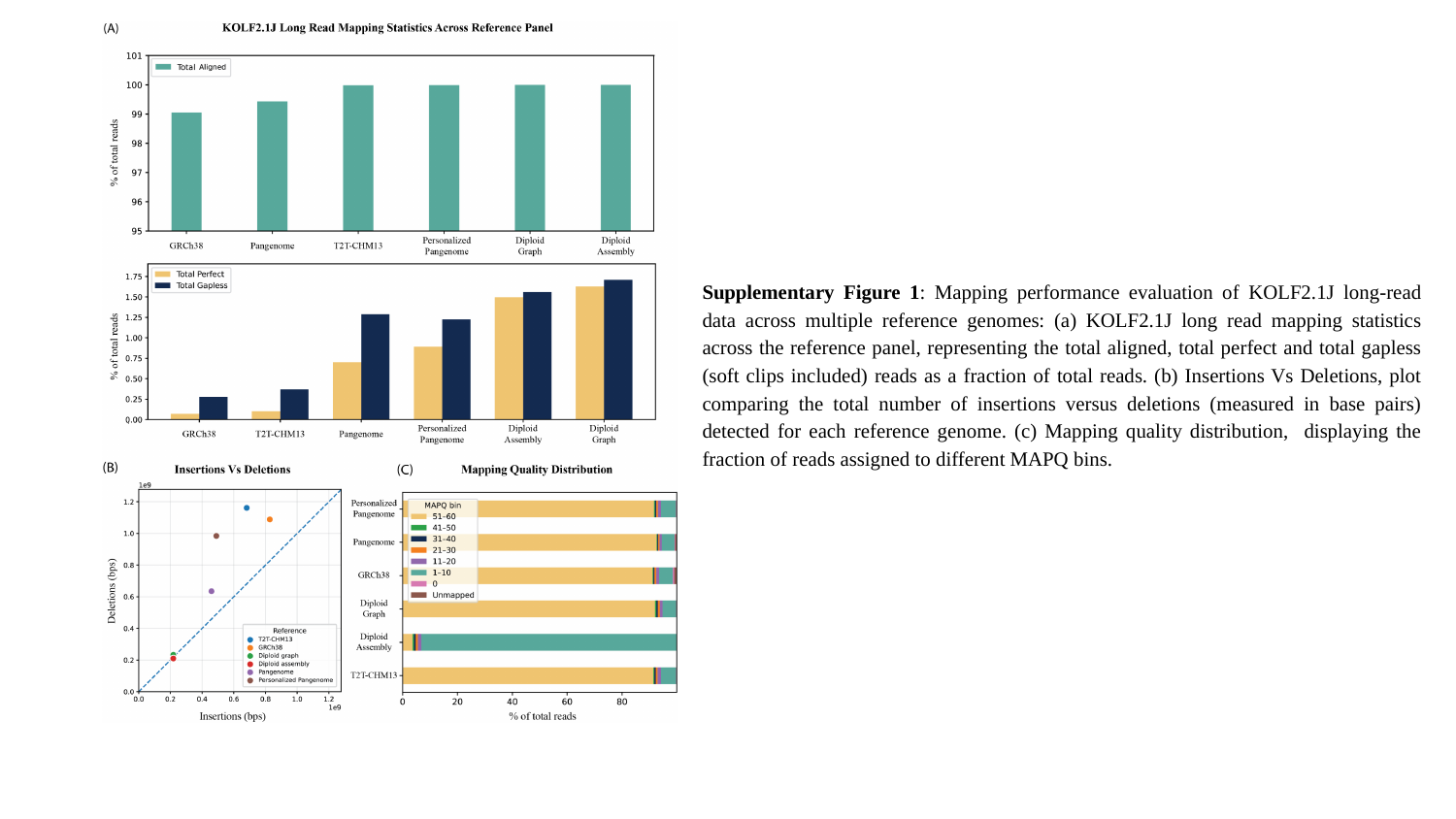

Supplementary Figure 1: Mapping performance evaluation of KOLF2.1J long-read data across multiple reference genomes: (a) KOLF2.1J long read mapping statistics across the reference panel, representing the total aligned, total perfect and total gapless (soft clips included) reads as a fraction of total reads. (b) Insertions Vs Deletions, plot comparing the total number of insertions versus deletions (measured in base pairs) detected for each reference genome. (c) Mapping quality distribution, displaying the fraction of reads assigned to different MAPQ bins.
