## Supplementary Methods for "The complete genome of the KOLF2.1J reference iPSC line"

### Cell Differentiation Protocols

#### Astrocyte differentiation - UCL

Astrocytes were generated from neuronal precursors generated according to Shi et al. 2012, following and Ziff et al, 2025 to differentiate into astrocytes (Shi et al. 2012; Ziff et al. 2025) at a culture of 60-70 days. In brief, cells were maintained in N2B27 media, a 1:1 mixture of N2 (DMEM/F12 Glutamax, N2 supplement, non-essential amino acids, β-mercaptoethanol, penicillin/streptomycin, insulin) and B27 (Neurobasal, B27 supplement, L-glutamine) media. Glial progenitor expansion was supported in N2B27 supplemented with FGF2 (10 ng/mL), and cells were passaged using EDTA-mediated dissociation to selectively remove mature, postmitotic neurons. Proliferating glial progenitors were replated on Geltrex-coated flasks at 1:3–1:5 splits and fed 3–4 days post-passage; after approximately day 110, cells adopted a glial morphology. From day 150 (>15 passages), astrocytes were matured in N2B27 containing LIF (10 ng/mL) and BMP4 (10 ng/mL) for 2–4 weeks, with media changes twice weekly.

#### Astrocyte and oligodendrocytes differentiation - NYSCF

Astrocytes and oligodendrocytes were differentiated using a previously established protocol that generates glial-enriched cultures (Douvaras and Fossati 2015a) All reagents and steps of the differentiation are extensively described in previous publications (Douvaras and Fossati 2015b; Barbar et al. 2020a)(Clayton et al. 2024; Prakash et al. 2024; Douvaras et al. 2014). Briefly, OLIG2+ glial progenitors were generated with dual smad inhibition plus patterning with smoothened agonist and retinoic acid. This is done to recapitulate in vitro the pMN domain where oligodendrocytes originate during fetal development. Cultures were grown in monolayer until day 12, when cells were lifted with a cell scraper to allow aggregation into OLIG2+ spheres. Spheres were then plated around day 30 into poly-ornithine/laminin coated wells at a density of 20-30 per well of a 6-well plate. As spheres attached to the well, progenitors started crawling out and filled the entire well within a couple of weeks. During this time, cultures were fed with a medium that contains growth factors and other molecules to promote gliogenesis and oligodendrocyte commitment, such as PDGF (10 ng/ml), IGF-1 (10ng/ml), NT3 (10ng/ml), Triiodothyronine (T3, 60ng/ml), dbCaMP (1µM), biotin (100ng/ml), and HGF (5ng/ml). At the end of the differentiation, astrocytes were isolated through cell sorting using the CD49f surface marker (Barbar et al. 2020b). The CD49f negative fraction is enriched for oligodendrocyte progenitor cells and oligodendrocytes, but still contains some neurons and early progenitors. Cell pellets for RNA or DNA extraction were obtained directly after CD49f sort by transferring the cell suspensions into eppendorf tubes and centrifugation. Cell pellets were flash frozen and stored at -80˚C.

#### Microglia differentiation

Microglia were differentiated as reported in Brownjohn et al 2018 (Brownjohn et al. 2018). This protocol followed an established method for the derivation of primitive macrophage precursors (PMPs) as a starting point for microglia differentiation (Karlsson et al. 2008; van Wilgenburg et al. 2013). In brief, iPSCs were passaged to single cells with TrypLE Express (Gibco, 12604013) and plated in 100uL embryoid body medium (EBM) (10 μM ROCK inhibitor, 50 ng/mL BMP-4, 20 ng/mL SCF, and 50 ng/mL VEGF-121 in E8). The following day, 80 µL of medium was removed from each well and replaced with 100 µL of EBM without ROCK inhibitor. Embryoid bodies were cultured for an additional 2 days, with daily media changes using fresh EBM without ROCK inhibitor. Then, ~200 embryoid bodies were transferred into a T75 flask and cultured in 20 mL hematopoietic medium (2 mM GlutaMax, 100 U/mL penicillin, 100 μg/mL streptomycin, 55 μM β-mercaptoethanol, 100 ng/mL M-CSF, and 25 ng/mL IL-3 in X-VIVO 15 (Lonza, LZBE02-060F), after which medium was exchanged every 2 days. Microglia progenitor cells (MPCs) typically appeared between Day 12 and Day 18. The cells were collected by centrifugation, and approximately 4 million cells per flask were seeded into non-coated 75 cm² flasks for final differentiation in Microglia Maturation Media (MMM), consisting of Advanced RPMI, 2 mM GlutaMax™, 100 ng/mL IL-34, and 10 ng/mL GM-CSF. Cells were further differentiated for 10 additional days, with full MMM media changes every other day.

#### Forebrain neuron differentiation

Cells were differentiated into forebrain neurons using a dual-SMAD inhibition protocol initially described by Burkhardt et al (Burkhardt et al. 2013). In short, iPSCs were grown on Matrigel-coated plates in E8 media (ThermoFisher, A1517001) until 90% confluent, after which they were transitioned to neuronal differentiation (N3) media (50% DMEM/F12, 50% Neurobasal™ with 1× penicillin-streptomycin, 0.5× B-27™ minus vitamin A, 0.5× N2 supplement, 1× GlutaMAX™, 1× NEAA, 0.055 mM 2-mercaptoethanol and 1 μg/ml Insulin) plus 1.5 μM dorsomorphin (Tocris Bioscience, 3093) and 10 μM SB431542 (Stemgent, 04-0010). Media was replaced every day for 11 days and cells were fed with additional N3 media without dorsomorphin and SB431542 until day 16. On differentiation days 16-20 N3 media was supplemented with 0.05μM retinoic acid. On day 20, cells were split and were then fed daily with N4 media (same as N3 plus 0.05 μM retinoic acid, 2 ng/ml BDNF and 2 ng/ml GDNF) until day 26 when they were pelleted for DNA and RNA isolation.

#### Transcription factor-NGN2 differentiation

The human NGN2 under a tetracycline-inducible promoter was expressed in KOLF2.1J iPSCs as previously described (Fernandopulle et al. 2018) using the PiggyBac system for delivery (Cookson and Flores 2023). Subsequently, iPSCs were differentiated into neurons as previously described (Pantazis et al. 2022) with some modifications (Flores et al. 2023). Briefly, the KOLF2.1J cells were transfected with PB-TO-oNGN2 vector (Addgene #198397) and EFa1-Transposase (Addgene plasmid #172116) at a 1:3 ratio (transposase:vector) and a total of 3µg of DNA mix, using Lipofectamine Stem Transfection Reagent (Thermo Fisher Cat. # STEM00008). This was followed by selection after 24-48 hours with 1 to 8 µg/ml of puromycin (Sigma-Aldrich Cat. #P9620) for a minimum of 72 hours. The selected NGN2-iPSCs were singularized using Accutase (Thermo Fisher Cat. # A111050) and plated at a density of 1.6x10^6^ cells into a Matrigel-coated (1:100, Corning Cat. # 354277) 100 mm dish with Neuronal Induction Media (NIM) containing DMEM/F12-HEPES (Thermo Fisher Cat. #. 11330032), 1X N2 Supplement (Thermo Fisher Cat. # 17502048), 1X Non-Essential Amino Acids (Thermo Fisher Cat. # 11140-050), 1X Glutamax (Thermo Fisher Cat. # 35050-061), 50 nM Chroman 1 (MedChemExpress Cat. # HY-15392) and 2 µg/mL Doxycycline (Sigma-Aldrich Cat. # D9891). NIM was replaced every day for 3 more days. Immature neurons were dissociated on day 4 using Accutase and replated at a density of 8x10^6^ cells into a poly-L-ornithine coated (PLO, 0.1 mg/ml, Sigma-Aldrich Cat. # P3655) 100 mm dish using Neuronal Maturation Media (NMM) for day 4 and day 7, containing, DMEM/F12-HEPES: Brainphys Neuronal Medium (1:1) (Stem Cell Technologies Cat. # 5790), 1X N21-MAX Supplement (R&D Systems Cat. # AR008), 10 ng/mL GDNF (PeproTech Cat. # 450-10), 10 ng/mL BDNF (PeproTech Cat. # 450-02), 10 ng/mL NT3 (PeproTech Cat. # 450-03), 1 µg/mL Laminin (R&D Systems Cat. # 3446-005-01), 2 µg/mL Doxycycline, 5-Fluoro-2′-deoxyuridine 1 µM (FdU, Sigma-Aldrich Cat. # F0503) and Uridine (Sigma-Aldrich Cat. # U3003). For day 10 to day 28 medium was replaced using NMM containing Brainphys Neuronal Media only plus all the supplements used at day 4 and 7. For neuronal maintenance after day 4, half-medium changes were performed every 3-4 days until day 28.

### Extraction Protocols

#### High Molecular Weight DNA Extraction

Full extraction and processing information for our cell high molecular weight (HMW) DNA can be found on protocols.io (Billingsley et al. 2022). In short, DNA from KOLF2.1J iPSCs was extracted manually from 1-5x10^6 million frozen cell pellets using the Nanobind Tissue Big DNA kit (Circulomics/PacBio). Samples were then size-selected with PacBio’s Short Read Eliminator Kit *(SS-100-101-01)* to deplete DNA fragments up to 25kb. The size selected DNA was then sheared to a target size of 30kb on the Diagenode Megaruptor3 using the Fluid + needles at speed 45. For all samples, DNA length was assessed by running 1ul the TapeStation 4200 (Agilent) or through a Fluoroskan (Thermo Fisher). DNA concentration was assessed using the dsDNA BR assay (Invitrogen Q32850) on a Qubit fluorometer (Thermo Fisher). This process was then repeated for the rest of the differentiated cell states.

#### Ultra-Long DNA extraction

Ultra high molecular weight DNA (UHMW) was extracted from KOLF2.1J iPSC using the Nanobind CBB Big DNA Kit and the UHMW DNA Aux Kit (PacBio) following the PacBio UHMW DNA Nanobind Extraction protocol (no longer available). The extracted UHMW DNA was then taken straight into library preparation for sequencing.

#### RNA Extraction

RNA was extracted from KOLF2.1J frozen 5x10^6 cell pellets following Thermo Fisher’s TRIzol User Guide (Pub. No. MAN0001271). In short, 0.75mL of TRIzol Reagent was added to each cell pellet after which cells were lysed and homogenized. 0.15mL of chloroform was added to the homogenized sample and centrifuged at 12,000 x *g* for 15 minutes at 4C to separate layers. The upper aqueous phase was removed and taken forward to RNA isolation. Here, 0.375mL of 100% isopropanol was added and the sample was centrifuged at 12,000 x *g* for 15 minutes at 4C, and the supernatant was subsequently removed. The RNA pellet was then washed with 75% ethanol and resuspended in RNAse-free water. RNA quantification was performed on Agilent’s 4200 Tapestation with the RNA Screentape. All RNA extractions resulted in a RIN ~9.

### Data Generation

#### Oxford Nanopore DNA Library Preparation and Sequencing

Extracted and processed HWM DNA from KOLF2.1J in multiple cell states was prepared for sequencing using Oxford Nanopore’s SQK-LSK110 library preparation kit following protocol modifications described on protocols.io (Billingsley et al. 2022). These modifications included a higher DNA input amount (4.5 ug), the use of short fragment buffer (SFB) during adapter ligation and clean up, and a longer final elution time (20 minutes at 37C). 400 ng of prepared library per sample was then loaded on to R9.4.1 PromethION flow cells and sequenced for 72 hours using the Minknow software. During these 72 hours, each sample underwent a wash and reload based on SQK-LSK110 instructions when pore occupancy dropped to ~2000 pores, leading to a total of 2-3 loads per flow cell.

Extracted UHMW DNA from KOLF iPSC underwent library preparation with the Oxford Nanopore SQK-ULK001 Kit and PacBio Nanobind Ultra Long Library Preparation Kit (NB-900-601-01) (no longer available). 75uL of the prepared library was then loaded and sequenced on R9.4.1 PromethION flow cells for 72 hours using the Minknow software. The same wash and reload guidelines as above were followed.

#### PacBio DNA Library Preparation and Sequencing

Samples were sent to Psomagen (Rockville, Maryland) for PacBio HiFi sequencing. Per Psomagen, HMW DNA was quantified using the Quant-iT PicoGreen dsDNA Assay Kit (cat. #P7589, Thermo Fisher Scientific) on a VictorX2 multilabel plate reader (PerkinElmer). DNA integrity and size distribution were assessed using the Femto gDNA 165kb Analysis kit (Part Number: FP-1002-0275) on the Agilent Femto Pulse System (Part Number: M5330AA) and DNA purity was assessed by NanoDrop’s 260/280 and 260/230 ratios. Libraries were prepared using the SMRTbell Prep Kit 3.0 (PacBio, cat. #102- 182-700) according to the manufacturer’s procedure. HMW genomic DNA was first evaluated on the Agilent Femto Pulse system to ensure quality (GQN10 kb ≥7.0). Short fragments were removed with the Short Read Eliminator (SRE), after which DNA was sheared using the Megaruptor 3 system (Diagenode, B06010003) with shear speed 29~31 to achieve a target insert size of 15–18 kb. Post-shearing cleanup was performed with SMRTbell cleanup beads, followed by DNA damage repair, A-tailing, adapter ligation, nuclease treatment, and AMPure PB bead size selection. The final SMRTbell libraries were annealed with sequencing primers, bound to polymerase (Revio SPRQ™ chemistry), and purified prior to sequencing.

PacBio HiFi sequencing was performed on the Revio system (PacBio) using SMRT Cells loaded with polymerase-bound SMRTbell libraries (prepared with the SMRTbell Prep Kit 3.0). The sequencing was performed using the Revio SPRQ polymerase kit (103-520-100), Revio SPRQ sequencing plate (103-504-900), and Revio SMRT Cell tray (102-202-200). Prepared libraries were diluted to the recommended concentration prior to sequencing. Sequencing plates and SMRT Cells were loaded onto the Revio instrument according to the manufacturer’s protocol. Data collection and real-time monitoring were performed through SMRT Link.

#### Oxford Nanopore RNA Library Preparation and Sequencing

200ng of total RNA from KOLF2.1J in multiple cell states was prepared for sequencing using Oxford Nanopore’s cDNA-PCR SQK-PCS111 library preparation kit and protocol with modifications. These modifications included an additional bead clean up after reverse transcription with 11.25 uL of RNAse-free XP beads, SFB washes, and elution into 22.5uL of elution buffer. The PCR settings during amplification were adjusted to set annealing to 12 cycles. After library preparation, total RNA was quantified using the Qubit dsDNA HS Assay Kit (Invitrogen, Q32851). 22 fmol, or 20 ng of prepared library, was loaded on to R9.4.1 PromethION flow cells and sequenced for 72 hours using the Minknow software.

#### PacBio RNA Library Preparation and Sequencing

RNA quality and quantity were assessed using the Agilent TapeStation 4200 system with RNA ScreenTape (Cat #5067-5576, Agilent Technologies) and RNA ScreenTape Reagents (Cat #5067-5577, Agilent Technologies), together with the Quant-iT™ RiboGreen RNA Assay Kit (Cat #R11490, Thermo Fisher Scientific), according to the manufacturers’ protocols.

For full-length cDNA synthesis, 300 ng of total RNA per sample was processed using the Iso-Seq Express Kit 2.0 (Cat #103-071-500, Pacific Biosciences). The resulting cDNA was then used to generate full-length RNA libraries using the Kinnex Full-Length RNA Library Prep Kit (Cat #103-072-000, Pacific Biosciences). The libraries were individually barcoded and pooled.

Sequencing was performed on the PacBio Revio System over three runs, each utilizing one Revio SMRT Cell (Cat #102-202-200, Pacific Biosciences), the Revio SPRQ Polymerase Kit (Cat #103-520-100, Pacific Biosciences), and the Revio SPRQ Sequencing Plate (Cat #103-504-900, Pacific Biosciences), following the manufacturer’s recommended protocol.

#### Hi-C Library Preparation and Sequencing

Hi-C libraries were generated from 10x10^6 iPSC cells using the Arima High Coverage Hi-C+ kit (Arima Genomics A201030) and the Arima Library Prep Module kit v2 (Arima Genomics A303011), following the manufacturer's protocols. The library was then sequenced on an Illumina HiSeq X system to achieve approximately 600M reads.

#### CAGE-seq library preparation and sequencing

To check the presence of 5′ caps and identify transcription start sites, we generated CAGE sequencing data for three of our cell types (iPSC, neuron, and microglia). CAGE library prep and sequencing was performed by DNAFORM (<https://www.dnaform.jp/en/>). In brief, RNA quality was assessed with a Bioanalyzer (Agilent) to ensure all lines had a RIN > 8. First-strand cDNAs were transcribed to the 5′ ends of capped RNAs and attached to CAGE ‘barcode’ tags. The samples were then sequenced on an Illumina NextSeq 500 and the sequenced CAGE tags were mapped to the human hg38 genome using BWA software (version 0.5.9).

### Custom Mapping Read Alignment and Parameter Optimization

To perform mapping experiments, we first built indexes with vg autoindex. For linear targets (T2T-CHM13, GRCh38, and the diploid assembly), we indexed the reference FASTA. For graph targets, we indexed the corresponding GFA or GBZ. Reads were then mapped with vg giraffe using presets appropriate for the sequencing data type. For personalized pangenome graph mapping, we first counted read k-mers with KMC and provided the resulting .hapl and .kff files during alignment.

Although vg giraffe is optimized for pangenome graphs, we also applied it to linear references and benchmarked performance against BWA-MEM2. Using KOLF short reads aligned to T2T-CHM13 with default parameters, vg giraffe produced ~8 million fewer aligned reads overall and ~5 million fewer perfectly aligned reads than BWA-MEM2. This difference is attributed to vg giraffe’s default --hard-hit-cap (500), which filters out reads with highly frequent minimizers, whereas BWA-MEM2 retains these reads, usually with low mapping quality scores.

Raising --hard-hit-cap to 9000 increased both the total number of aligned reads and the number of perfectly aligned reads, reducing the residual perfectly aligned reads difference to ~0.5 million. Further increases in --hard-hit-cap could recover additional reads, but at increased computational cost. After this adjustment, BWA-MEM2 still reported ~3 million additional aligned reads relative to vg giraffe; however, these reads were predominantly supported by low-confidence alignments that are filtered by vg giraffe under its scoring and mapping-quality criteria. Accordingly, all vg giraffe mapping results reported in this study were generated with --hard-hit-cap 9000.

Barbar, Lilianne, Tanya Jain, Matthew Zimmer, et al. 2020a. “CD49f Is a Novel Marker of Functional and Reactive Human iPSC-Derived Astrocytes.” *Neuron* 107 (3): 436–453.e12.

Barbar, Lilianne, Tanya Jain, Matthew Zimmer, et al. 2020b. “CD49f Is a Novel Marker of Functional and Reactive Human iPSC-Derived Astrocytes.” *Neuron* 107 (3): 436–453.e12.

Billingsley, Kimberley J., Pilar Alvarez Jerez, Abigail Miano-Burkhardt, and Cornelis Blauwendraat. 2022. *Processing Frozen Human Blood Samples for Population-Scale Oxford Nanopore Long-Read DNA Sequencing SOP*. August 22. <https://www.protocols.io/view/processing-frozen-human-blood-samples-for-populati-b6fhrbj6.pdf>.

Brownjohn, Philip W., James Smith, Ravi Solanki, et al. 2018. “Functional Studies of Missense TREM2 Mutations in Human Stem Cell-Derived Microglia.” *Stem Cell Reports* 10 (4): 1294–1307.

Burkhardt, Matthew F., Fernando J. Martinez, Sarah Wright, et al. 2013. “A Cellular Model for Sporadic ALS Using Patient-Derived Induced Pluripotent Stem Cells.” *Molecular and Cellular Neurosciences* 56 (September): 355–364.

Clayton, Benjamin L. L., Lilianne Barbar, Maria Sapar, et al. 2024. “Patient iPSC Models Reveal Glia-Intrinsic Phenotypes in Multiple Sclerosis.” *Cell Stem Cell* 31 (11): 1701–1713.e8.

Cookson, Mark, and Erika Lara Flores. 2023. *iNDI PiggyBac-TO-hNGN2 Transfection Protocol Version 1*. August 8. <https://www.protocols.io/view/indi-piggybac-to-hngn2-transfection-protocol-versi-cyd6xs9e.pdf>.

Douvaras, Panagiotis, and Valentina Fossati. 2015a. “Generation and Isolation of Oligodendrocyte Progenitor Cells from Human Pluripotent Stem Cells.” *Nature Protocols* 10 (8): 1143–1154.

Douvaras, Panagiotis, and Valentina Fossati. 2015b. “Generation and Isolation of Oligodendrocyte Progenitor Cells from Human Pluripotent Stem Cells.” *Nature Protocols* 10 (8): 1143–1154.

Douvaras, Panagiotis, Jing Wang, Matthew Zimmer, et al. 2014. “Efficient Generation of Myelinating Oligodendrocytes from Primary Progressive Multiple Sclerosis Patients by Induced Pluripotent Stem Cells.” *Stem Cell Reports* 3 (2): 250–259.

Fernandopulle, Michael S., Ryan Prestil, Christopher Grunseich, Chao Wang, Li Gan, and Michael E. Ward. 2018. “Transcription Factor-Mediated Differentiation of Human iPSCs into Neurons.” *Current Protocols in Cell Biology* 79 (1): e51.

Flores, Erika Lara, Andy Qi, Luke Reilly, Marianita Santiana, Michael Ward, and Mark Cookson. 2023. *iNDI Transcription Factor-NGN2 Differentiation of Human iPSC into Cortical Neurons Version 2*. June 6. <https://www.protocols.io/view/indi-transcription-factor-ngn2-differentiation-of-bzchp2t6.pdf>.

Karlsson, Karl R., Sally Cowley, Fernando O. Martinez, Michael Shaw, Stephen L. Minger, and William James. 2008. “Homogeneous Monocytes and Macrophages from Human Embryonic Stem Cells Following Coculture-Free Differentiation in M-CSF and IL-3.” *Experimental Hematology* 36 (9): 1167–1175.

Pantazis, Caroline B., Andrian Yang, Erika Lara, et al. 2022. “A Reference Human Induced Pluripotent Stem Cell Line for Large-Scale Collaborative Studies.” *Cell Stem Cell* 29 (12): 1685–1702.e22.

Prakash, Priya, Hediye Erdjument-Bromage, Michael R. O’Dea, et al. 2024. “Proteomic Profiling of Interferon-Responsive Reactive Astrocytes in Rodent and Human.” *Glia* 72 (3): 625–642.

Shi, Yichen, Peter Kirwan, and Frederick J. Livesey. 2012. “Directed Differentiation of Human Pluripotent Stem Cells to Cerebral Cortex Neurons and Neural Networks.” *Nature Protocols* 7 (10): 1836–1846.

Wilgenburg, Bonnie van, Cathy Browne, Jane Vowles, and Sally A. Cowley. 2013. “Efficient, Long Term Production of Monocyte-Derived Macrophages from Human Pluripotent Stem Cells under Partly-Defined and Fully-Defined Conditions.” *PloS One* 8 (8): e71098.

Ziff, Oliver J., Gustavo Morrone Parfitt, Sarah Jolly, et al. 2025. “Mutations in PSEN1 Predispose Inflammation in an Astrocyte Model of Familial Alzheimer’s Disease through Disrupted Regulated Intramembrane Proteolysis.” *Molecular Neurodegeneration* 20 (1): 73.
